# Programmable DNA cleavage by Ago nucleases from mesophilic bacteria *Clostridium butyricum* and *Limnothrix rosea*

**DOI:** 10.1101/558684

**Authors:** Anton Kuzmenko, Denis Yudin, Sergei Ryazansky, Andrey Kulbachinskiy, Alexei A. Aravin

## Abstract

Argonaute (Ago) proteins are the key players in RNA interference in eukaryotes, where they function as RNA-guided RNA endonucleases. Prokaryotic Argonautes (pAgos) are much more diverse than their eukaryotic counterparts but their cellular functions and mechanisms of action remain largely unknown. Some pAgos were shown to use small DNA guides for endonucleolytic cleave of complementary DNA *in vitro*. However, previously studied pAgos from thermophilic prokaryotes function at elevated temperatures which limits their potential use as a tool in genomic applications. Here, we describe two pAgos from mesophilic bacteria, *Clostridium butyricum* (CbAgo) and *Limnothrix rosea* (LrAgo), that act as DNA-guided DNA nucleases at physiological temperatures. In contrast to previously studied pAgos, CbAgo and LrAgo can use not only 5’-phosphorylated but also 5’-hydroxyl DNA guides, with diminished precision of target cleavage. Both LrAgo and CbAgo can tolerate guide/target mismatches in the seed region, but are sensitive to mismatches in the 3’-guide region. CbAgo is highly active under a wide range of conditions and can be used for programmable endonucleolytic cleavage of both single-stranded and double-stranded DNA substrates at moderate temperatures. The biochemical characterization of mesophilic pAgo proteins paths the way for their use for DNA manipulations both *in vitro* and *in vivo*.

## INTRODUCTION

Argonaute proteins are an integral part of the eukaryotic RNA interference machinery. They bind small noncoding RNAs and utilize them for guided cleavage of complementary RNA targets or indirect gene silencing by recruiting additional factors (1–3). Argonaute proteins are also found in bacterial and archaeal genomes (4–6). Structural and biochemical studies of a few prokaryotic Ago (pAgo) proteins – predominantly from thermophilic bacterial and archaeal species – showed that they can function as endonucleases *in vitro* (7–11) and may provide cell defense against foreign genetic elements *in vivo* (8,12,13). pAgos are thus proposed to act as a bacterial system of innate immunity acting against invasive genetic elements (5,6,12–15).

Prokaryotic Argonaute proteins can be classified into several clades, including long pAgos (further subdivided into two clades, long-A and long-B), short pAgos and PIWI-RE proteins (4,6,16). All pAgos characterized so far belong to the long pAgo clade and include the N, PAZ, MID and PIWI domains (except for AfAgo that has lost its N and PAZ domains), also present in eAgos (Fig. 1A). The MID and PAZ domains are responsible for binding of the 5’ and 3’ ends of a guide nucleic acid molecule, respectively (7,17–21). In contrast to eAgos that use exclusively small RNA guides, the majority of characterized pAgos bind single-stranded DNA (ssDNA) guides, including AaAgo (from *Aquifex aeolicus*) (10), AfAgo (*Archaeoglobus fulgidus*) (22), TtAgo (*Thermus thermophilus*) (13), PfAgo (*Pyrococcus furiosus*) (8), and MjAgo (*Methanocaldococcus jannaschii*) (21). RsAgo (*Rhodobacter sphaeroides*) (12) and MpAgo (*Marinitoga piezophila*) (7) were shown to bind small RNA guides. The specificity towards RNA or DNA guides cannot be inferred from pAgo sequence and has to be determined experimentally. Binding of the guide to pAgo changes its conformation to expose nucleotides in the so-called seed region (2-8 nt from the guide 5’-end) in solution in an A-form helix to facilitate target recognition (7,20,21). While eAgos universally recognize complementary RNA as a target, studies of a few long pAgos suggested that they employ their guides to bind DNA targets (7,8,11,12,19,22,23). Complementary interactions between the guide and double-stranded DNA (dsDNA) target require unwinding of the DNA helix; however, pAgos lack intrinsic helicase activity, so the molecular mechanism of this step remains to be understood.

**Figure 1.**
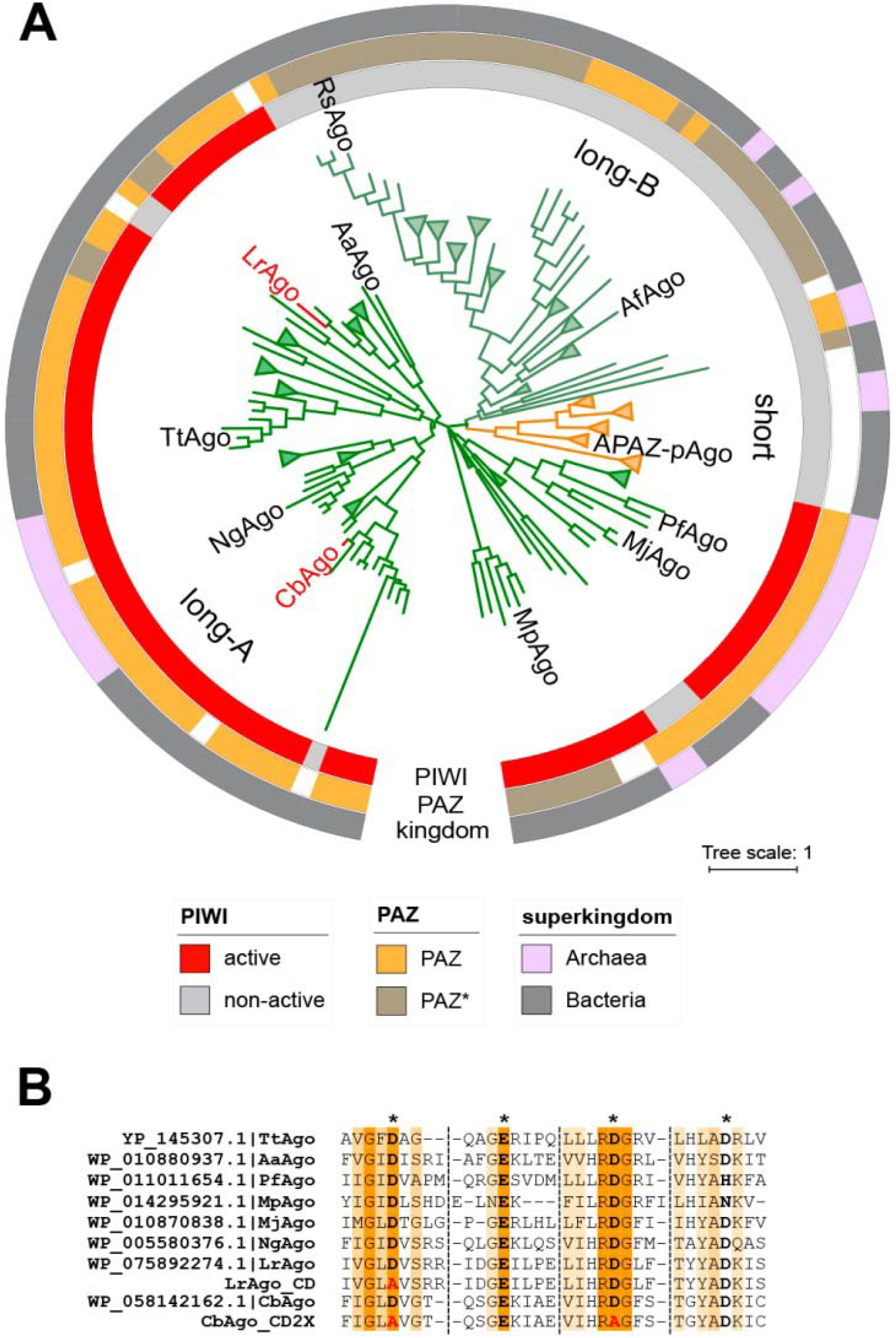
CbAgo and LrAgo are mesophilic pAgo proteins with an intact catalytic tetrad. (A) The circular phylogenetic tree of nonredundant set of pAgos constructed based on multiple alignment of the MID-PIWI domains (4). The pAgo proteins were annotated as follows (from the inner to the outer circles): the type of the PIWI domain, depending on the presence of the catalytic tetrad DEDX; the presence and the type of the PAZ domain (full-length PAZ, ochre; incomplete PAZ, brownish; no PAZ, white); the superkingdom to which the corresponding pAgo belongs. Long-A, Long-B, and short-pAgo clades are indicated. The scale bar represents the evolutionary rate calculated under the JTT+CAT evolutionary model (4). The positions of biochemically characterized pAgos are shown; CbAgo and LrAgo are shown in red. (B) Multiple sequence alignment of conserved amino acid residues (marked by color and asterisks) of the DEDX tetrad localized in the PIWI domain of pAgos. The catalytically dead variants of the of the LrAgo and CbAgo proteins (LrAgo_CD and LrAgo_CD2X) with substituted aminoacid substitutions within the catalytic tetrad are also shown. Rs, *Rhodobacter sphaeroides*; Af, *Archaeoglobus fulgidus*; Pf, *Pyrococcus furiosus*; Mj, *Methanocaldococcus jannaschii*; Mp, *Marinitoga piezophila*; Cb, *Clostridium butirycum; Ng, Natronobacterium gregoryi*; Tt, *Thermus thermophilus*; *Lr, Limnothrix rosea;* Aa, *Aquifex aeolicus.*

The PIWI domain of most long-A pAgos has an RNase H-like fold with the active site containing the DEDX (X = N, D or H) catalytic tetrad, which endows these proteins with endonuclease activity (4,6). Dissociation of the processed target strand after cleavage was shown to be the rate-limiting step in catalysis, which could be overcome at increased temperatures in the case of pAgos from thermophilic prokaryotes (19,24,25). Some pAgos such as RsAgo have substitutions in the catalytic tetrad making them deficient in endonuclease activity (12). The active site of the PIWI domain slices the complementary target at a single site, between the tenth (10’) and eleventh (11’) nucleotides starting from the guide 5’-end (7–10,13). In contrast to the Cas9 protein that has two distinct endonuclease domains allowing it to cut both strands of dsDNA upon target recognition, pAgos have only one active site, so only one DNA strand can be cleaved by a single complex. Mismatches between the guide and target molecules in the seed region decrease the target binding rate by previously studied pAgos and perturb the slicer activity, while mismatches adjacent to the cleavage site abolish it completely (9,17,24,26). In addition to guide-dependent processing of ssDNA targets, several thermophilic pAgos were shown to perform slow guide-independent cleavage of dsDNA termed chopping (8,11,27). Short DNAs generated by the chopping activity of pAgos might then associate with pAgos and be used as guides for further specific target cleavage. Chopping was proposed to facilitate the onset of immunity against foreign DNA (11,27).

pAgos are programmable endonucleases that may potentially be used as a tool for DNA manipulations *in vitro* and *in vivo*, including molecular cloning and genome editing applications (14,16,28). However, catalytically active pAgos characterized to date were derived from thermophilic species and function at elevated temperatures. Furthermore, it is not clear if pAgos are able to process dsDNA substrates at moderate temperature without additional partners due to their inability to unwind dsDNA. These concerns have cast doubt on the practicality of pAgos as a tool for DNA manipulation, *e.g.* (29,30). Here we describe two Argonaute proteins from mesophilic prokaryotes, bacillus *Clostridium butyricum* (CbAgo) and cyanobacterium *Limnothrix rosea* (LrAgo). We show that both pAgos are DNA-guided DNA nucleases that function at much lower temperatures than other pAgos studied to date. We characterize activities of LrAgo and CbAgo under a wide range of conditions and reveal functional differences in the efficiency and fidelity of DNA processing by the two proteins. Finally, we demonstrate that CbAgo is able to perform precise guide-dependent cleavage of dsDNA when supplied with two guides targeting both strands of dsDNA target. Finding of programmable pAgo endonucleases that are able to process dsDNA targets at moderate temperatures opens the way for their use as a tool in DNA technology.

## MATERIALS AND METHODS

### Protein expression and purification

Nucleotide sequences of CbAgo (WP_058142162.1; *C. butyricum* strain TK520) and LrAgo (WP_075892274.1; *L. rosea* strain IAM M-220) were codon-optimized using IDT Codon Optimization Tool for expression in *E. coli*, the genes were synthesized by IDT core facility and cloned into p-ET28b expression vectors in frame with the N-terminal His_6_ tag. Catalytically dead mutants were obtained by site-directed mutagenesis using QuikChange Lightning Multi mutagenesis kit (Agilent). Expression was carried out as described in ref. (31) with minor modifications. Briefly, *E. coli* BL21(DE3) cells carrying the expression plasmid were adapted to the high ionic strength conditions by overnight cultivation in the LBN medium at 37°C. The cells were transferred into fresh LBN (1:500 inoculation) supplemented with 1 mM bethaine and aerated at 37°C until OD_600_ 0.6. At this point, the cultures were cooled down to 18°C, induced with 0.25 mM IPTG and aerated for 12 hours at 18°C. The cells were collected by centrifugation and stored at −80°C for further protein purification.

Both wild-type and mutant proteins were purified by the same three-step scheme. Cell pellet was resuspended in Ni-NTA chromatography buffer A (50 mM Tris-HCl, 0.5 M NaCl, 20 mM imidazole, 5% glycerol, 1 mM TCEP pH 7.5) supplemented with EDTA-free protease inhibitor cocktail (Roche) and disrupted on a high-pressure homogenizer at 18000 psi. The lysate was clarified by centrifugation at 32000 g for 30 min and the supernatant was loaded onto HisTrap HP column (GE Healthcare). The column was washed extensively with buffer A, then with buffer A containing 60 mM imidazole, and the proteins were eluted with buffer A containing 300 mM imidazole. Fractions containing pAgos were concentrated by ultrafiltration using Amicon 50K filter unit (Millipore) and purified on the Superose 6 10/300 GL column (GE Healthcare) equilibrated with buffer GF (10 mM HEPES-NaOH, 0.5 M NaCl, 5% glycerol, 1 mM DTT pH 7.0). Fractions containing pAgos were pulled and loaded onto the Heparin FF column (GE Healthcare) equilibrated with buffer GF, washed with at least 10 column volumes of the same buffer and eluted with a linear NaCl gradient (0.5 - 1 M). Samples containing CbAgo and LrAgo (both eluted at 650-700 mM) were aliquoted and flash-frozen in liquid nitrogen. The purity of the final protein samples was assessed by denaturing PAGE followed by silver staining. The protein concentration was determined by Qubit protein assay kit (Thermo Fischer Scientific).

### Nucleic acid cleavage assays

Most cleavage assays were performed at 5:2:1 Ago:guide:target molar ratio at 37°C, unless otherwise indicated. 500 nM CbAgo or LrAgo was mixed with 500 nM guide DNA in reaction buffer containing 25 mM Tris-HCl pH 7.4, 100 mM NaCl, 5% glycerol and 5 mM MnCl_2_, and incubated at 37°C for 10 min for guide loading. Target DNA was added to 100 nM. The reaction was stopped after indicated time intervals by mixing the samples with equal volumes of stop solution (8 M urea, 20 mM EDTA, 0.005% Bromophenol Blue, 0.005% Xylene Cyanol). The cleavage products were resolved by 19% denaturing PAGE, stained with SYBR Gold and visualized with Typhoon FLA 9500 or ChemiDoc XP. In experiment shown on the Fig.2C all reactions were incubated at indicated temperatures simultaneously using a PCR thermocycler (T100, Bio-Rad). In plasmid cleavage assays, 2 nM of plasmid pSRKKm_t was added to the reaction mixtures, and the reactions were stopped by treatment with Proteinase K at 4°C. The samples were mixed with 6xPurple Loading Dye (New England Biolabs) and cleavage products were resolved on 1% native agarose gels.

### EMSA assays

Reaction mixtures containing the required components (pAgo, guides and targets) were incubated in the reaction buffer at 37°C for 10 min, then mixed with 5x loading dye (5x TBE, 50% glycerol) and resolved by 10% native PAGE buffered with TBE at 4°C. Nucleic acids were stained with SYBR Gold (Invitrogen) and visualized using Typhoon FLA 9500.

### *K*_d_ measurements

Apparent dissociation constants (*K*_d_) for guide DNA binding were determined using double-filter assay as described (32). Briefly, 10 pM 5’-[P^32^]-labeled oligonucleotide was mixed with increasing amounts of CbAgo or LrAgo protein in 50 μl of binding buffer (10 mM HEPES-NaOH, 100 mM NaCl, 5 % glycerol, 100 μg/ml BSA, 5 mM MnCl_2_, pH 7.0) and incubated at 37°C for 40 min. The sampes were filtered through nitrocellulose membrane (Merck-Millipore) layered on top of nylon Hybond N+ (GE Healthcare) equilibrated with the ice-cold binding buffer. After three washing steps with 200 μl of the same buffer the membranes were removed, air-dried and analyzed by phosphorimaging, using the ImageQuant (GE healthcare) and Prism 8 (GraphPad) software. The data were better fitted with the Hill equation with a Hill coefficient of 2-2.5 rather than with a standard binding isotherm (p < 0.0001), indicative of some cooperativity in guide binding (possibly reflecting pAgo interactions with both 5’-and 3’-guide ends).

For the competition binding assay, increasing amounts of unlabeled 5’-OH or 5’-P guide DNA was added to the 50 μl reaction mixture containing 100 pM of radiolabeled 5’-P DNA guide, followed by the addition of 100 pM LrAgo protein. After 1 hour incubation at 37°C, the samples were processed as described above. The data were fitted using the one-site competitive binding model.

## RESULTS

### CbAgo and LrAgo use small DNA guides for endonucleolytic cleavage of ssDNA targets at ambient temperature

Many members of the Ago family in both pro- and eukaryotes rely on the RNaseH-like active site in their PIWI domain for endonucleolytic cleavage of nucleic acid targets, while others have substitutions in the catalytic tetrad which likely make them inactive as nucleases (see Introduction). Based on our phylogenetic analysis of prokaryotic Ago proteins (4), we selected two pAgos from mesophilic bacteria, cyanobacterium *L. rosea* (LrAgo) and anaerobic bacillus *C. butyricum* (CbAgo), belonging to different clades of long pAgos (Fig. 1A). Sequence alignments showed that both LrAgo and CbAgo have four conserved catalytic resides (DEDD) in the PIWI domain suggesting that they potentially have catalytic endonuclease activity (Fig. 1B).

To study biochemical properties of CbAgo and LrAgo we expressed and purified each protein. Codon-optimized genes encoding both proteins were chemically synthesized and cloned. In addition to the wild-type proteins, we obtained their catalytically inactive variants with substitutions of one or two out of four catalytic tetrad residues (D541A/D611A in CbAgo and D516A in LrAgo). The proteins were expressed in *E. coli* cells and purified using sequential Ni-NTA-affinity, size-exclusion and cation-exchange chromatography steps (see Fig. S1 and Materials and Methods for details). Examination of purified pAgos showed high purity of the samples with a single band of the expected molecular weight (Fig. S1).

We first studied the preference of CbAgo and LrAgo for RNA and DNA guides and targets using guide-dependent target cleavage assay (Fig. 2A, Fig. S2A). CbAgo and LrAgo were loaded with 18 nt 5’-phosphorylated DNA or RNA guides followed by the addition of complementary 50 nt long single-stranded DNA or RNA targets (Fig. 2B, Fig. S2, Table S1). After incubation at 37°C in buffer with divalent metal ions (Mn^2+^ in most experiments) the products were resolved on denaturing gel. In reactions containing guide (g) DNA and target (t) DNA, we observed guide-dependent target cleavage at a single site between target positions 10’ and 11’ (Fig. 2C, Fig. S2). No cleavage products were observed in the absence of pAgo proteins (Fig. 2C). The cleavage required the presence of intact catalytic tetrad in the PIWI domain as point mutations in the tetrad eliminated the activity of both CbAgo and LrAgo (Fig. S2B, Fig. S3B). Remarkably, CbAgo showed comparable levels of endonuclease cleavage between 30 and 54 °C and still retained some activity at 60 °C (Fig. 2D). LrAgo was less active than CbAgo at most temperatures, but was stimulated at 50-54 °C, and inactivated at 60 °C (Fig. 2D). Thus, CbAgo and LrAgo are active at ambient conditions and are sufficiently stable to perform this reaction at elevated temperatures.

**Figure 2.**
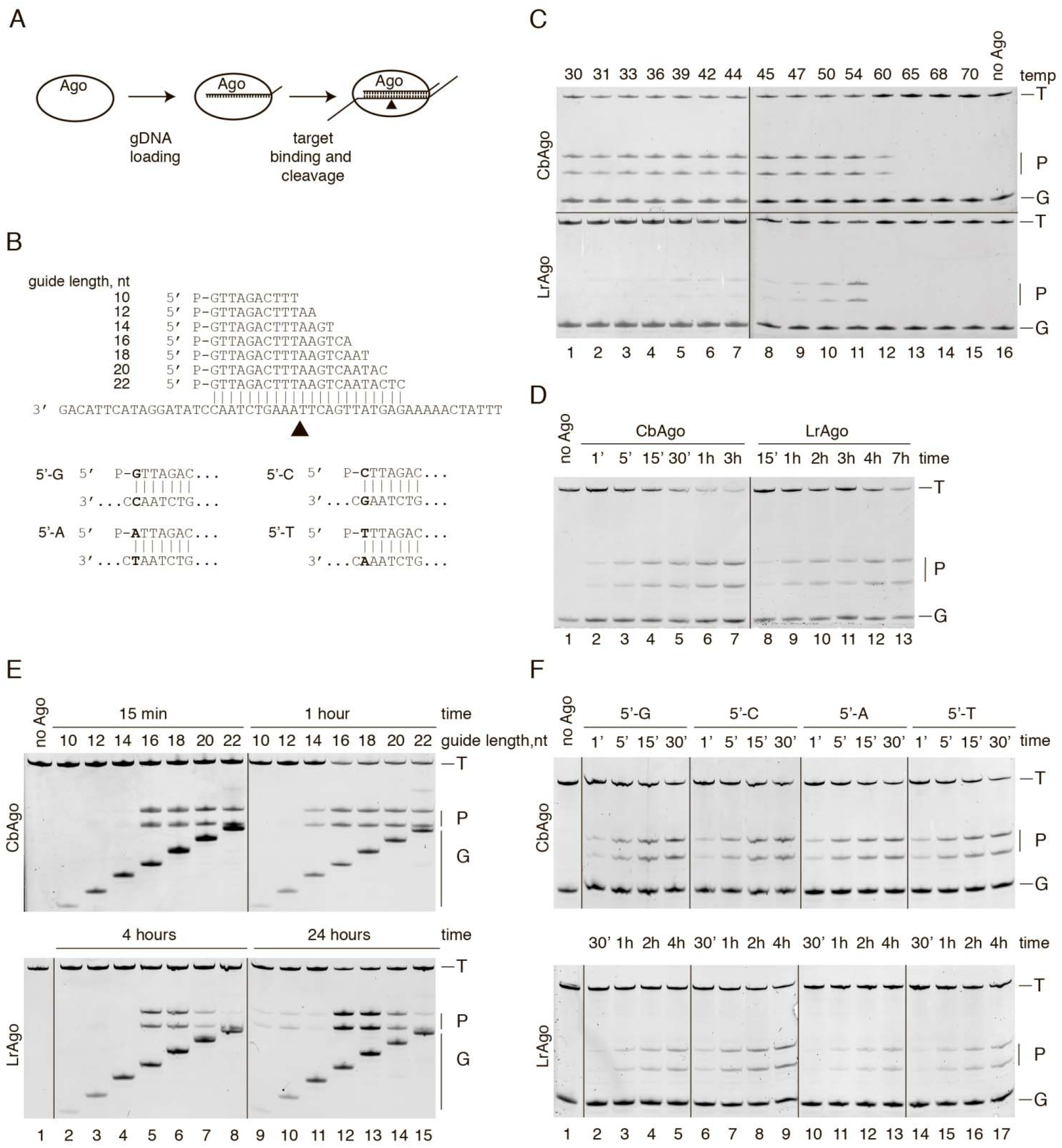
CbAgo and LrAgo are DNA-dependent DNA-endonucleases. (A) Scheme of the *in vitro* assay. (B) Guide DNAs and target DNAs used in the experiments. Black triangle indicates the predicted cleavage site. For guide variants with different 5′-ends only 5’-parts of the corresponding guide-target pairs are shown. (C) CbAgo and LrAgo are active at moderate temperatures. The assay was performed at the 5:5:1 pAgo:guide:target molar ratio for 30 min at indicated temperatures. CbAgo is active under a broad temperature range, while LrAgo activity peaks at 54°C. Moreover, CbAgo is more stable compare to LrAgo at elevated temperatures. (D) Time-course analysis of ssDNA cleavage by CbAgo and LrAgo at 37°C. The reactions were performed at the 5:2:1 pAgo:guide:target molar ratio for indicated time intervals. (E) Cleavage assay with guide DNAs of varied length. In case of LrAgo some cleavage is observed even with the shortest guides (10-14 nt) upon prolonged incubation. (E) Preferences for the 5’-guide nucleotide. LrAgo shows a slight preference for 5’-G and 5’-C, while CbAgo slices ssDNA with the same efficiency with all four guide variants. T, target; P, cleavage products; G, guide.

No substrate cleavage was observed with other combinations of guide and target molecules (*i.e.*, gRNA-tDNA, gDNA-tRNA and tRNA-gRNA) (Fig. S2). Analysis of nucleic acids co-purified with CbAgo and LrAgo from *E. coli* cells also demonstrated that both proteins bind small DNAs *in vivo* (unpublished observations). Thus, similarly to the majority of previously characterized pAgos, CbAgo and LrAgo act as DNA-guided DNA nucleases, but unlike them are active at moderate temperatures.

To further explore the catalytic properties of CbAgo and LrAgo, we measured the kinetics of target cleavage in reactions containing 18 nt gDNA and complementary 50 nt tDNA at 37°C (Fig. 2D). We found that CbAgo can cleave the ssDNA target significantly faster than LrAgo, with cleavage products detected already after 1 min, while for LrAgo comparable cleavage was observed after 15 minutes (Fig. 2C). Thus, we used longer incubation times for LrAgo in most experiments described below. We next analyzed the effects of divalent cations and other reaction conditions on guide-dependent DNA cleavage. Both CbAgo and LrAgo had the highest endonuclease activity in the presence of Mn^2+^ and are much less active with Mg^2+^ (only marginal activity was observed for LrAgo). LrAgo was inactive in the presence of Co^2+^, Cu^2+^ or Zn^2+^, while CbAgo could also catalyze target cleavage in the presence of Co^2+^ (Fig. S3B). Titration of Mn^2+^ ions showed that LrAgo activity peaks at 5 mM Mn^2+^ (Fig. S3C), which is significantly higher than concentrations optimal for Ago proteins characterized previously (e.g., 100 μM for PfAgo (8)). Mn^2+^ is an abundant cation in cyanobacterial cells (33), possibly explaining this unusual property of LrAgo. LrAgo is most active at NaCl concentrations of 50 to 100 mM but retains a certain level of cleavage at high ionic strength up to 750 mM NaCl (Fig. S3D). pH does not noticeably influence DNA cleavage by LrAgo within the tested range (6.8 – 8.0) (Fig. S3E).

Interestingly, we did not observe complete cleavage of the ssDNA substrate if the target was present in excess over the binary pAgo-guide complex, even after prolonged incubation at 37 °C (Fig. S4). This suggested that the reaction has a limited turnover, possibly due to slow dissociation of the product complex. At the same time, when we performed experiments with CbAgo at 55 °C, the efficiency of the reaction was significantly increased, suggesting that each binary guide-pAgo complex can process multiple target molecules under these conditions (Fig. S4).

Thus, in contrast to previously characterized pAgos from thermophilic species that are active only at high temperatures, CbAgo and LrAgo are mesophilic DNA-guided DNA nucleases that can potentially be active in eukaryotic cells at 37 °C.

### Determination of the optimal length and sequence of guide DNA

All eAgos and pAgos studied to date bind nucleic acid guides of specific length (34) (7,11–13). Furthermore, many eAgos and pAgos were also reported to have a bias for particular nucleotide residue at the first position of the guide molecule, which is accommodated in a protein pocket formed by the MID domain (12,18,21,35,36). To determine the role of guide DNA structure on pAgo-mediated target cleavage, we performed the cleavage reaction with guides of different lengths and sequences (see Table S1). First, we explored the role of the guide length by testing a series of DNA guides from 10 to 22 nt long that shared identical sequences at their 5’-ends, so that the predicted cleavage site was the same for all guides (Fig. 2B). 16-18 nt guides led to efficient target cleavage, while shorter guides diminished the reaction efficiency (Fig. 2E). However, even the shortest 10 and 12 nt guides were able to direct proper target cleavage by LrAgo (but not CbAgo), albeit with a significantly lower efficiency that required prolonged incubation (24 h, right panels in Fig. 2E). The cleavage efficiency for LrAgo was also significantly reduced with guides longer than 18 nt suggesting that extended duplex formation may prevent target cleavage (Fig. 2E).

To determine if CbAgo and LrAgo have a preference for the first nucleotide of the guide, we tested four guide variants with different 5′-terminal nucleotides but otherwise identical sequences (Fig. 2B). All four guides were able to direct cleavage of complementary targets by both CbAgo and LrAgo, with a small preference for 5′-G and 5′-C guides observed for LrAgo (Fig. 2F). Thus, the two pAgo proteins can likely direct cleavage of any desired sequence making them a flexible tool for DNA manipulation.

### CbAgo and LrAgo can use both 5′-P and 5’-OH guides for target cleavage, but with different precision

All eukaryotic and the majority of prokaryotic Ago proteins – including the best studied TtAgo and RsAgo – strongly prefer 5′-phosphorylated nucleic acid guides due to multiple contacts formed between the phosphate group and amino acid residues in the 5′-end binding pocket of the Ago MID domain (17,18,20,21,35–37). However, pAgo from *M. piezophila* (MpAgo) has a unique 5′-end binding pocket that confers it the ability to bind 5′-OH guides and exclude 5′-phosphorylated molecules (7). Analysis of the 5′-binding pocket in the MID domain of LrAgo and CbAgo revealed substitutions of two out of six amino acid residues in the specific motif involved in interactions with the Me^2+^ ion and the guide 5′-phosphate relative to TtAgo (Fig. 3A).

**Figure 3.**
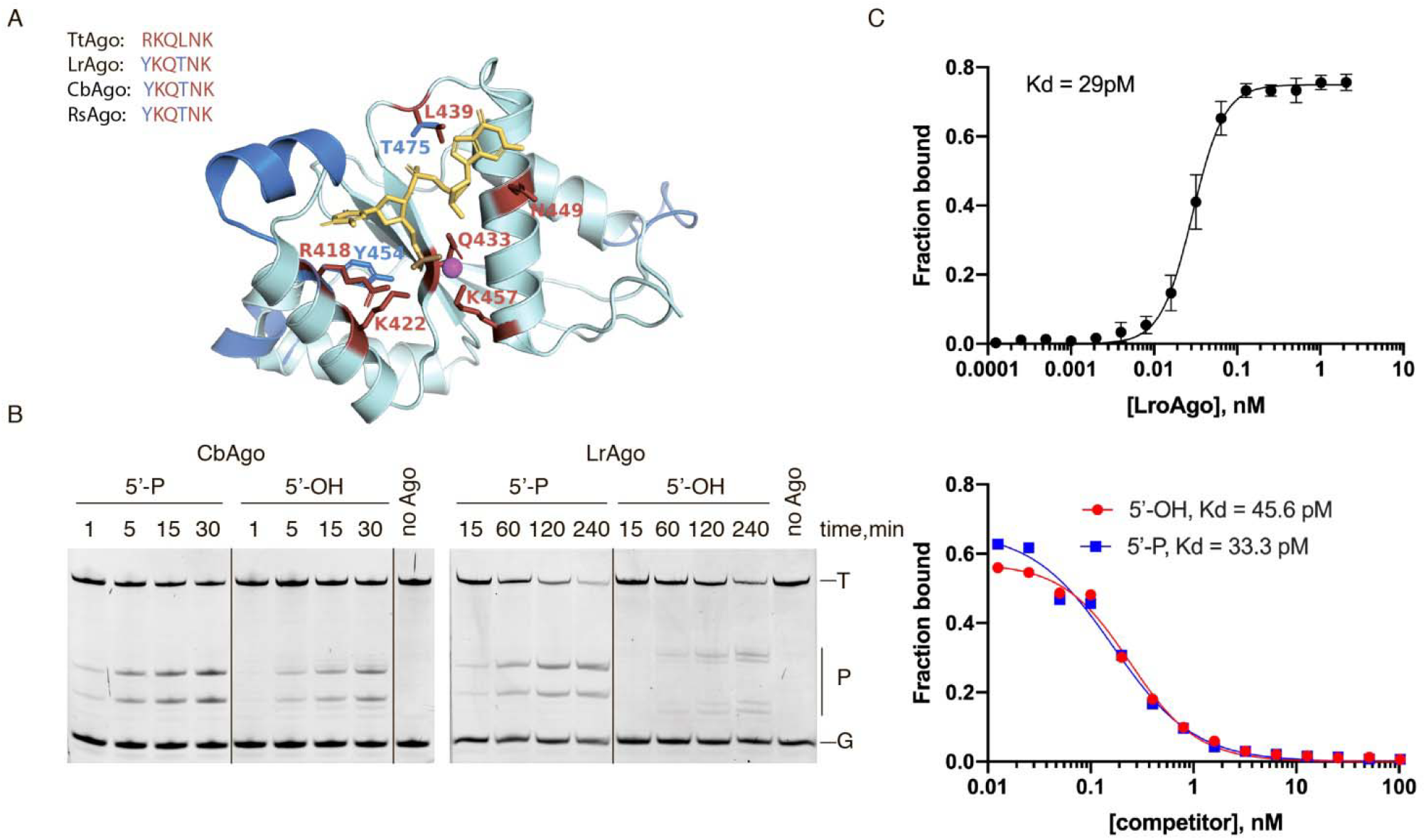
CbAgo and LrAgo can utilize both 5’-phosphorylated and 5’-hydroxyl DNA guides at 37°C. (A) A 3D model of the LrAgo MID domain aligned to the corresponding structure of the TtAgo MID domain in complex with guide DNA (PDB: 3HO1) was built using a SWISS-MODEL portal. Conserved amino acid residues of the YKQTNK motif (red) and the Mg^2+^ ion (magenta) involved in the binding of the first two guide nucleotides (yellow) are highlighted. Elements of secondary structure and amino residues specific to LroAgo are shown in blue. The sequences of the conserved MID motif involved in interactions with the guide 5’-end in various pAgos are shown above the model. (B) Programmable ssDNA cleavage by CbAgo and LrAgo in the presence of either 5’-P or 5’-OH guides at 37°C. The reactions were performed at the 5:2:1 pAgo:guide:target molar ratio for indicated time intervals. Notice the shift in the slicing position in the case of nonphosphorylated guide with LrAgo. (C) Binding of 18 nt phosphorylated DNA guide by LrAgo at 37°C. The fraction of bound DNA was plotted against protein concentration and fitted using a model of specific binding with the Hill slope. The corresponding *K*_d_ value is indicated on the graph (95 % confidence interval is 27.4 to 30.7 pM). (D) Competition binding assay. Radiolabeled guide DNA was combined with increasing amounts of unlabeled 5’-P (blue) or 5’-OH (red) competitor, and incubated with LrAgo at 37°C. The data are plotted as a fraction of bound DNA against competitor concentration; corresponding *K*_d_ values are indicated (95 % confidence intervals are 25.1 – 43.9 pM for the 5’-P guide and 33.7 – 61.7 pM for the 5’-OH guide). Means and standard deviations from 3 independent experiments are shown.

We studied the ability of CbAgo and LrAgo to use 5’-P and 5’-OH guides in the cleavage reaction. Surprisingly, both CbAgo and LrAgo could use 5’-OH DNA guides to cleave the target DNA, although the rate of the reaction was somewhat lower compared to 5’-P guides of identical sequence (Fig. 3B). Interestingly, when LrAgo was loaded with the 5’-OH DNA guide, the target cleavage was observed 1-2 nucleotides upstream of the canonical site, between target positions 8’-9’ and 9’-10’ relative to the guide 5′-end (Fig. 3B). The ability of pAgos to use 5’-OH guides to cleave target ssDNA was lost at 55°C (Fig. S5), suggesting that interactions with the 5’-phosphate are required to stabilize the binary pAgo-guide complex at elevated temperature.

Since LrAgo revealed unexpected changes in the cleavage site with the 5’-OH guide, we further measured the equilibrium dissociation constants (*K*_d_) for guide binding using a filter-based titration assay (Fig. 3C). LrAgo associated with 5′-phosphorylated guides with an apparent *K*_d_ value of 29 pM demonstrating a much higher affinity to nucleic acid guides compared to other studied pAgos (see Discussion). CbAgo also revealed exceptionally high affinity to 5’-P guides, which even exceeded that of LrAgo (*K*_d_ of 5-10 pM). To compare the ability of LrAgo to bind 5′-P and 5′-OH guides we used a competition assay, in which LrAgo was incubated with a mixture of 5′-P^32^-labeled guide and increasing amounts of unlabeled guide of the same sequence either containing or lacking the 5’-phosphate. Surprisingly, LrAgo revealed essentially the same affinities towards both types of DNA guides in this assay (Fig. 3C). Electrophoretic mobility shift assay (EMSA) also showed that the 5’-OH and 5’-P guides form binary complexes with LrAgo with comparable affinities (Fig. S6). Taken together, the results demonstrate that CbAgo and LrAgo are able to bind both 5’-P and 5’-OH DNA guides and use them for target DNA cleavage. The nature of the 5’-guide group is also critical for defining the exact position of the cleavage site by LrAgo. In contrast, the majority of Ago proteins are thought to have strong preference for 5′-phosphorylated guides, while MpAgo was shown to have strong preference for 5’-OH RNA guides.

### CbAgo and LrAgo tolerate mismatches in the seed region, but are sensitive to mismatches in the 3’-portion of guide DNA

Previous studies of eAgos and several pAgos, including AfAgo, TtAgo, RsAgo and MpAgo, showed that mismatches between the guide and target strands might have large effects on the efficiency of target cleavage (9,17,22,24,26,38–41). In particular, even a single mismatch in the seed region within 2-8 nts of the guide can lead to a significant decrease in the efficiency of target recognition and silencing for many eAgos as well as for pAgos (e.g. (9,17,18)). To study the effect of mismatches on target cleavage we designed a set of DNA guides, each containing a single mismatched nucleotide at a certain position (Table S1), and tested them in the cleavage reaction with CbAgo and LrAgo (Fig. 4).

**Figure 4.**
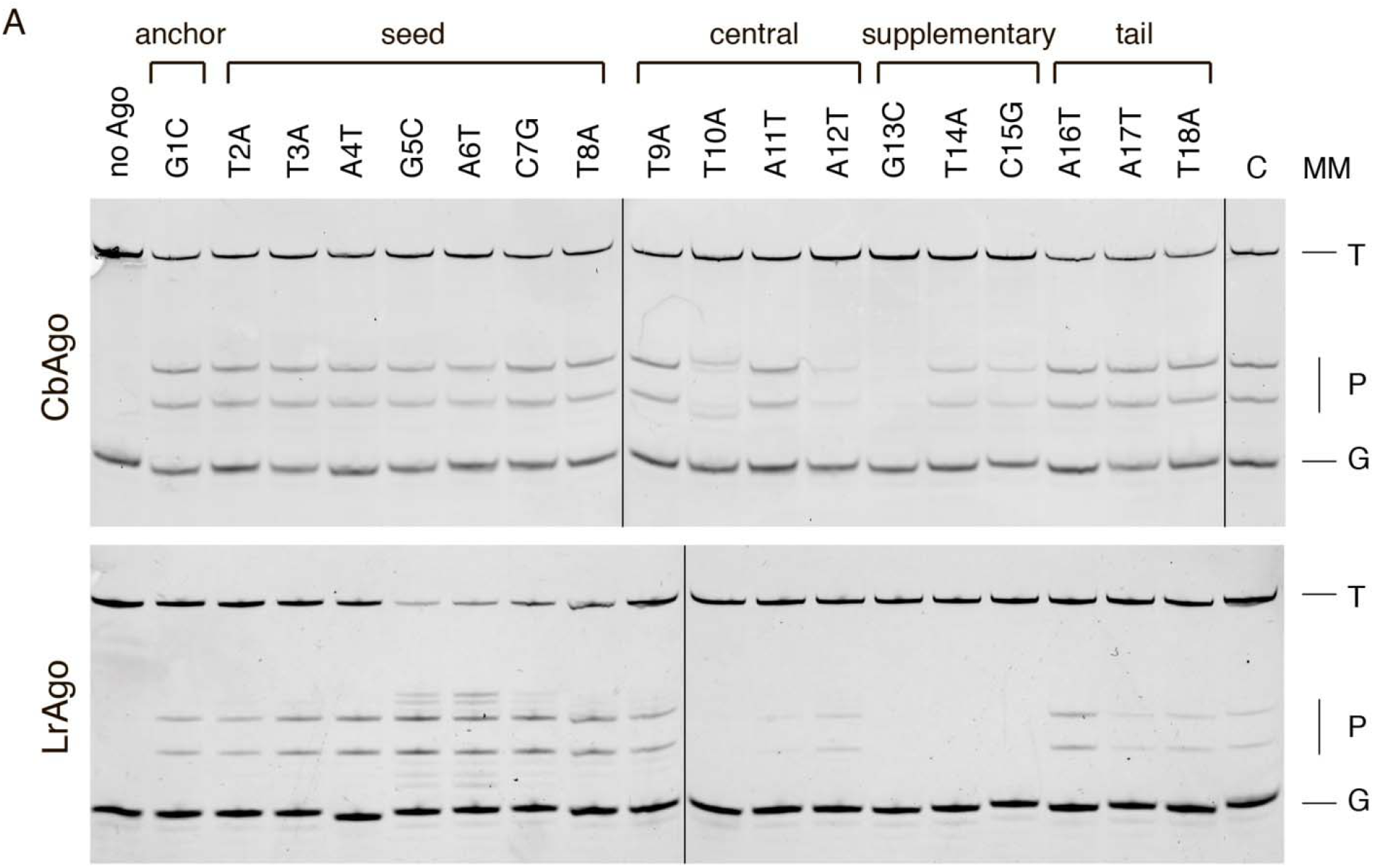
Effects of mismatches in the guide-target duplex on the slicing activity of CbAgo and LrAgo. (A) The assay was performed as in Fig. 3B except for the different Ago:guide:target molar ratio (5:5:1). Mismatch positions in different guide regions are designated above the gels; nucleotide substitutions relative to the standard guide sequence are indicated. Mismatches can cause a non-canonical shift in activity (*e.g.* position 10 for CbAgo, positions 5 and 6 for LrAgo), stimulate (seed region for LrAgo) or abolish (3’-supplementary region for both pAgos) the target cleavage. T, target; P, cleavage products; G, guide; C, control reactions with wild-type pAgos.

Surprisingly, mismatches in the seed region had little or no effect on the cleavage efficiency by CbAgo (Fig. 4A) and substantially increased target cleavage by LrAgo (Fig. 4B). Mismatches at positions 5 and 6 of the seed region also induced target cleavage at several additional sites located closer to the guide 5’-end (Fig. 4B). Mismatches downstream of the cleavage site, in the so-called supplementary region (guide positions 12-15), significantly decreased the efficiency of target cleavage by both CbAgo and LrAgo (Fig. 4A). Similarly, mismatches at the site of cleavage (10–11) led to a strong decrease in the target cleavage by LrAgo (Fig. 4B). However, in the case of CbAgo, mismatch at position 11 had no significant effect on cleavage, while mismatch at position 10 shifted the cleavage site one nucleotide closer to the guide 5’-end (Fig. 4A). Thus, in contrast to majority of Ago proteins studied to date, target cleavage by CbAgo and LrAgo is not inhibited and can be even stimulated by mismatches in the seed region but is decreased by mismatches in the 3’-supplementary guide region.

The effects of mismatches on the cleavage rate might be explained by changes in the formation of ternary pAgo:guide:target complex prior to cleavage or in the efficiency of endonucleolytic cleavage. To discriminate between these possibilities, we studied ternary complex formation by LrAgo, which is characterized by the lower rate of catalysis, using EMSA. In addition to the fully complementary guide-target pair, we tested mismatches at positions 4 and 11/14 that increased and decreased the rate of cleavage, respectively (see above). Almost all target DNA was bound by guide-loaded LrAgo within 10 minutes, regardless of the presence and position of the mismatched nucleotide (Fig. S7). Rapid binding of target DNA by LrAgo relative to the time required for its cleavage (Fig. 2D) suggests that target molecules reside within ternary complexes for extended time intervals prior to slicing. Overall, our data suggest that mismatches do not significantly affect the rate of target binding, but may instead change the rate of catalysis, possibly by changing the conformation of the ternary complex.

### Guide-free CbAgo and LrAgo can process plasmid DNA

While guide-loaded CbAgo and LrAgo can cut ssDNA substrates, the cleavage of dsDNA likely presents a bigger challenge for pAgos since the dsDNA duplex has to be unwound to form the ternary complex with guide-loaded pAgo. However, pAgos do not have helicase domains and, unlike the Cas9 protein, cannot perform DNA melting. Indeed, previous studies of TtAgo and PfAgo from thermophilic prokaryotes observed guide-dependent dsDNA cleavage only at elevated temperatures, which likely facilitated DNA melting (8,13). At the same time, thermophilic pAgos, TtAgo and MjAgo, were shown to process double-stranded plasmid DNA in a guide-independent manner (the so-called ‘chopping’), generating small DNA fragments that could be further loaded into pAgos and used as guides for subsequent target cleavage (11,27).

We tested the ability of CbAgo and LrAgo to cut supercoiled plasmid DNA at 37°C. The plasmid was incubated with empty (unloaded) pAgos, or with pAgos loaded with DNA guides designed to target the two DNA strands with predicted cleavage sites separated by 2 nt (Fig. 5A, B). Analysis of the cleavage products after incubation of the plasmid with empty LrAgo showed disappearance of the supercoiled (SC) form and increase in the open circular (OC) form likely containing single-stranded nicks in one or two strands (Fig. 5D). Furthermore, the linear form (LIN) and shorter DNA fragments that migrated on the gel as a light smear appeared upon prolonged incubation. No plasmid processing was observed with catalytically-dead (CD) LrAgo mutant with substitutions in the active site (Fig. 5D). The nicking and linearization of plasmid DNA was little affected by the addition of DNA guides, indicating that LrAgo likely cuts plasmid at random positions in a guide-independent manner. Therefore, although LrAgo demonstrates high accuracy in programmable slicing of ssDNA targets, it acts as a guide-independent endonuclease on double-stranded plasmid DNA.

**Figure 5.**
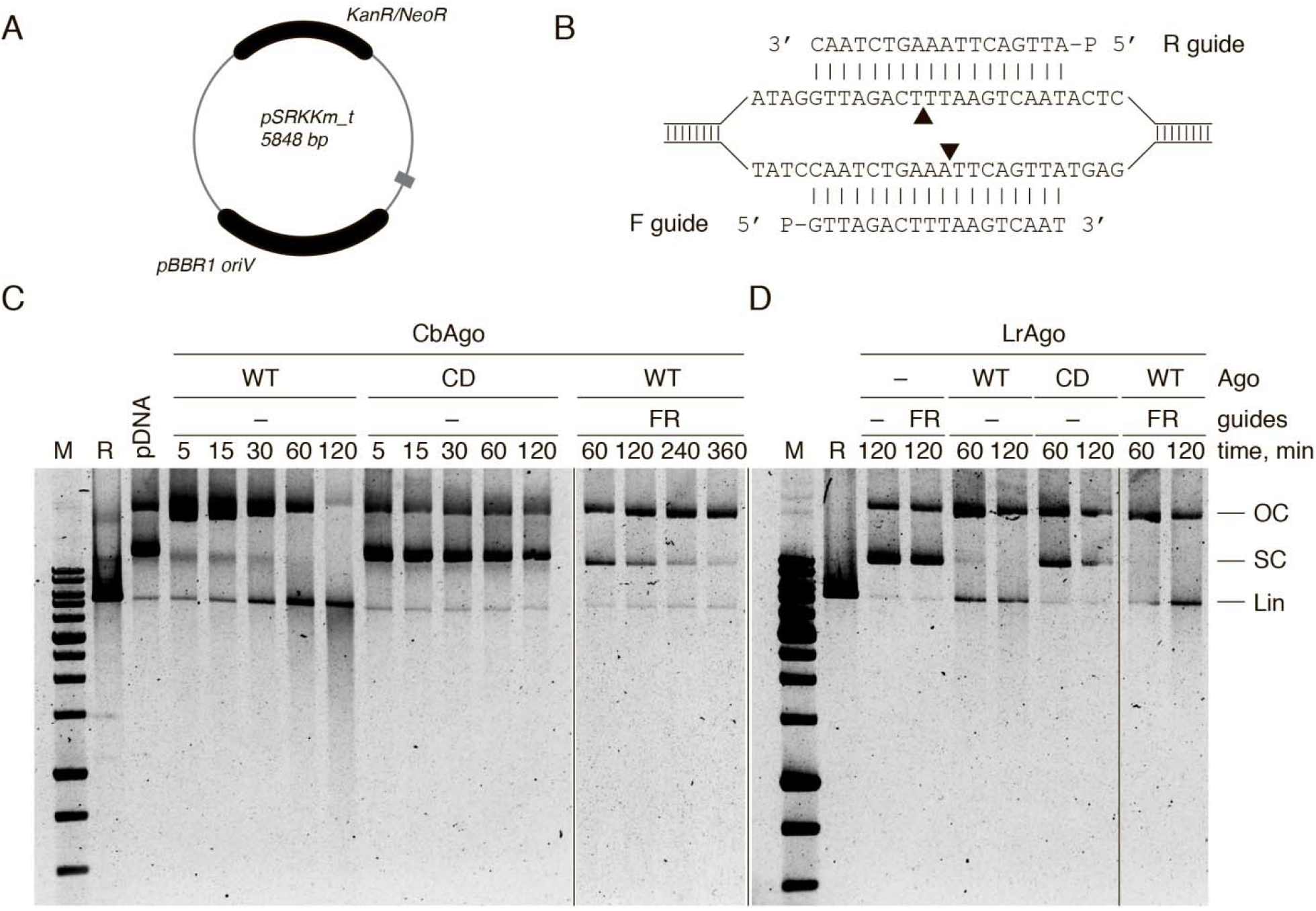
Guide-independent cleavage of plasmid DNA by CbAgo and LrAgo. (A) Scheme of the pSRKKm_t plasmid used in the assay. The target region is shown in grey. (B) Schematic representation of the target region with two DNA guides. Black triangles indicate the predicted cleavage sites. (C and D) pSRKKm cleavage by CbAgo (C) or LrAgo (D) in the presence or absence of DNA guides at 37°C. The reactions were carried out for the indicated time periods, resolved on the 1% agarose gel and stained with SYBR Gold. WT, wild-type; CD, catalytically dead CbAgo; FR, forward and reverse guide DNAs; M, molecular weight marker; R, control linear plasmid obtained after treatment with a restriction endonuclease; LIN, linearized plasmid; OC, open circular plasmid; SC, supercoiled plasmid.

Similarly to LrAgo, the wild-type, but not catalytically-dead, CbAgo could rapidly relax supercoiled plasmid in the absence of guide molecules, resulting in appearance of the open circular form (Fig. 5C, 5 min incubation time). This form gradually disappeared after prolonged incubation, accompanied by appearance of the linear form and a smear of shorter DNA products, indicative of the chopping activity (Fig. 5C, 120 min incubation). Unlike for LrAgo, the chopping activity of CbAgo was suppressed when it was loaded with the guides corresponding to the target plasmid (Fig. 5C, the ‘FR’ reaction). Thus, nonspecific processing of double-stranded DNA substrates is suppressed when CbAgo is bound to a guide molecule, which makes it a better candidate for targeted DNA cleavage.

### Guide-directed cleavage of double-stranded DNA by CbAgo

We further searched for conditions that would enhance the ability of CbAgo to use DNA guides for specific cleavage of dsDNA. Importantly, we observed no plasmid processing in the absence of guide molecules if the reaction was performed at 55°C (Fig. 6C). Thus, guide-independent chopping activity of CbAgo is suppressed at elevated temperature. The linear plasmid product was formed with high efficiency when the plasmid was incubated with CbAgo pre-loaded with two guide molecules targeting different strands of the plasmid at the same site (Fig. 6C). No chopping products of lower size were formed in this reaction. Next, we incubated the plasmid with CbAgo loaded with two pairs of guide molecules corresponding to different sites in the plasmid separated by ∼ 1 Kb (Fig. 6A). In these conditions, CbAgo cut plasmid DNA into two linear molecules with the size of fragments corresponding to sites targeted by guide DNAs (Fig. 6C). Therefore, CbAgo can perform dsDNA cleavage – likely by cutting each DNA strand independently of the other – with the precision similar to restriction endonucleases or the Cas9 nuclease, but without strict sequence requirements.

**Figure 6.**
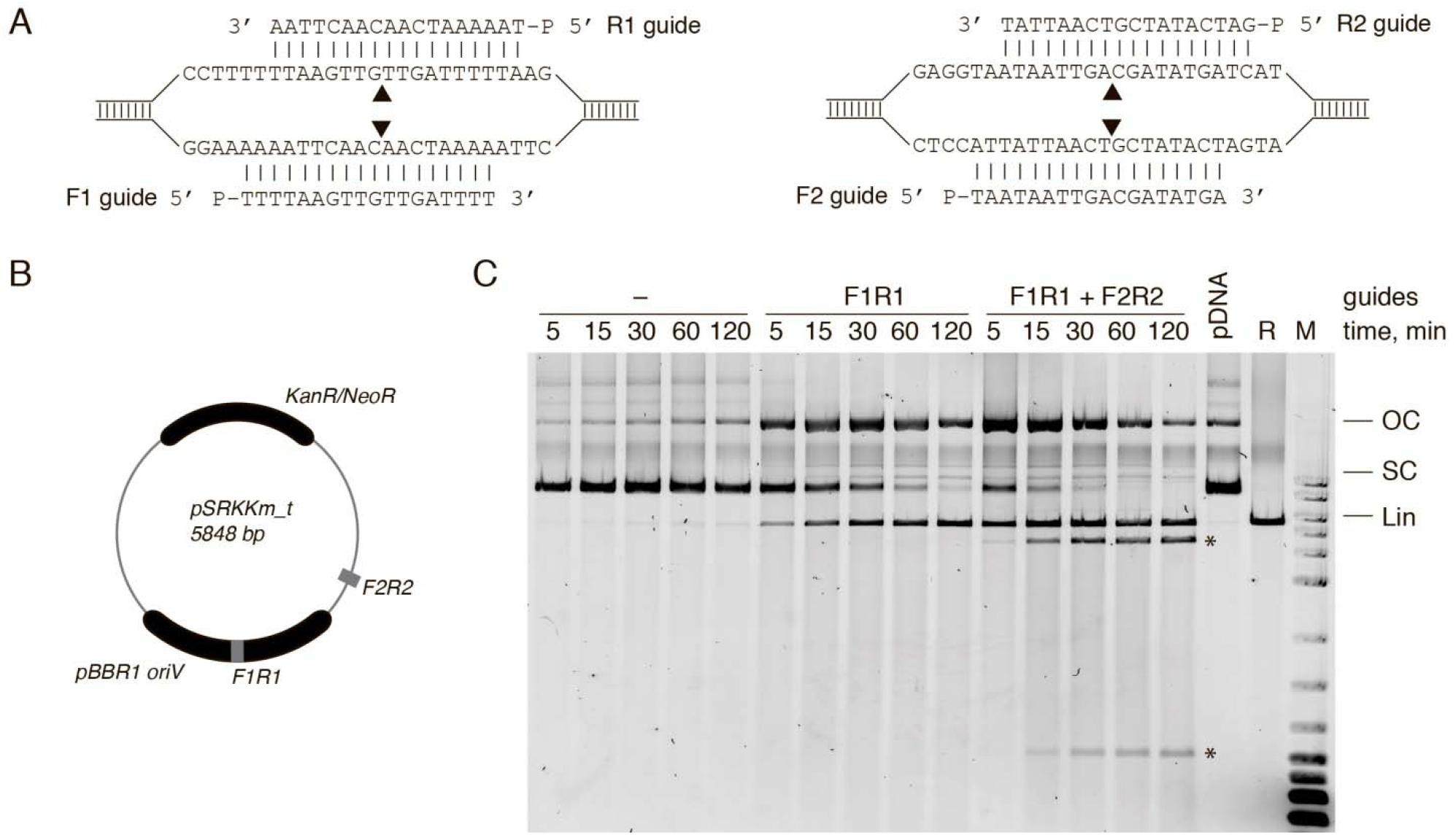
CbAgo acts as a programmable DNA-nuclease *in vitro*. (A) Schematic representation of the target regions of the pSRKKm_t plasmid with four DNA guides (F1R1 and F2R2). Black triangles indicate predicted cleavage sites. (B) Scheme of the pSRKKm_t plasmid with the two target regions shown in grey. (C) Plasmid cleavage by CbAgo with no guides, one pair of guides (F1R1) or two pairs of guides (F1R1 + F2R2) at 55°C. CbAgo programmed with four guides can efficiently excise the DNA region of interest (specific cleavage products are indicated with asterisks). All designations are the same as in Fig. 5.

## DISCUSSION

The majority of previously characterized pAgo proteins were derived from thermophilic bacterial and archaeal species and therefore are optimally active at high temperatures. In this study we present detailed characterization of pAgo proteins from mesophilic bacteria *C. butyricum* and *L. rosea*. We show that CbAgo and LrAgo are active endonucleases that can be programmed with small DNA guides to process target DNA substrates with high precision at moderate temperatures. Below, we compare the properties of CbAgo and LrAgo and suggest that they can be used for development of tools for manipulation of DNA *in vitro* and *in vivo*.

Both CbAgo and LrAgo are long catalytically active pAgo containing the complete DEDD catalytic tetrad in their PIWI domains. They perform precise DNA-guided slicing of ssDNA substrates *in vitro* at a wide range of temperatures (from 30 to 60 °C for CbAgo), and prefer Mn^2+^ as the catalytic ion. CbAgo is significantly faster than LrAgo at 37 °C and can act as an even more efficient multiple-turnover enzyme at elevated temperatures. Based on these and other properties described below, CbAgo appears a promising candidate for various genomic applications.

Both CbAgo and LrAgo can bind small DNA guides with exceptionally high affinities (with *K*_d_s in picomolar range), which markedly exceed those previously reported for other pAgos (*e.g.* ∼3 nM for MjAgo, ∼1 nM for RsAgo) (17,42). CbAgo and LrAgo show no obvious preference toward 5’-guide nucleotide during target cleavage indicating that both proteins can be programmed with DNA guides of any sequence permitting flexible choice of the target site. The efficiency of DNA cleavage by CbAgo and LrAgo is significantly reduced if guide length is below 16 nucleotides. However, slow target slicing could still be observed for LrAgo even with shorter guides (down to 10 nt) when the scissile bond (between target positions 10’ and 11’) was not flanked by base-paired nucleotides. TtAgo was also shown to use 9-10 nt DNA guides for target DNA cleavage, but this activity disappeared at increased temperatures (9). Thus, correct base-pairing around the cleavage position likely helps to stabilize target binding in the active site of pAgos.

In contrast to other studied Agos, both CbAgo and LrAgo are able to utilize 5’-OH guides for target cleavage with almost the same efficiency as 5’-P guides. The majority of eAgos and pAgos were reported to use 5’-P guides, and multiple contacts between the 5’-P group and the 5’-binding pocket in the MID domain are observed in the structures of several Ago-guide binary complexes (17,18,20,21,35–37). The notable exception is the RNA-guided MpAgo that binds exclusively to 5’-OH guides (7), and several other pAgos predicted to have a similar 5’-binding pocket (4,7). Interestingly, though eAgos universally associate with 5’-P guides *in vivo*, human Ago2 was shown to cut mRNA targets when bound to non-phosphorylated small RNA guides *in vitro* (43). This suggests that the ability of various Ago proteins to use non-phosphorylated guide may be underexplored.

Our recent bioinformatic analysis revealed several subtypes of the MID domain with substitutions of key residues involved in interactions with the 5′-group of guide molecule (4); most of them have not been characterized experimentally. Relative to TtAgo, LrAgo and CbAgo contain substitutions of two residues in the 5’-end binding pocket (Fig. 3A). However, these substitutions are also present in RsAgo that was shown to recognize 5’-phosphorylated guide molecules, similarly to TtAgo (Fig. 3A) (12). Furthermore, homology-based structural modeling suggests that interactions of the guide 5’-end with the MID pocket are overall very similar for TtAgo and LrAgo (Fig. 3A). Thus, additional protein-DNA interactions with other parts of the guide molecule may compensate for the loss of stabilizing interactions with the 5’-phosphate in LrAgo and CbAgo. Yet, interactions with the 5’-phosphate can stabilize the complexes under suboptimal conditions, such as increased temperature, as demonstrated for CbAgo that is not able to use 5’-OH guides to cut targets at 55°C.

In the case of LrAgo the cleavage site is shifted by 1-2 nucleotides upstream in the absence of the 5’-phosphate group in guide molecule. Changes in the slicing position were also observed for hAgo2 with nonphosphorylated guides (43). Such changes might be caused by sliding of the guide-target duplex in the active site in the absence of the 5’-P-MID interactions. Interestingly, MjAgo also demonstrated noncanonical target DNA cleavage 1-2 nucleotides closer to the guide 5’-end, suggesting that the guide-target duplex can be bound in several registers in the case of this pAgo (11,21,23). The role of the 5’-P-MID interactions in guide positioning in MjAgo remains to be established. Our results indicate that the guide 5’-phosphate can help to determine the correct register of the guide/target duplex relative to the active site of pAgo.

Previous studies of eAgos and pAgos demonstrated the importance of complementarity between the guide and the target for efficient repression: mismatches in the seed region (2-8 nt of the guide) reduce repression, while mismatches in the 3’-part (downstream of the cleavage site) are usually tolerated without significant loss of efficiency (9,17,22,24,26,38–41). In comparison with these studies, mismatches between the guide and target molecules have highly unusual effects on target cleavage by CbAgo and LrAgo. Mismatches in the seed region do not significantly affect (for CbAgo) or even increase (for LrAgo) the efficiency of target cleavage. Interestingly, recent analysis of zAgo2 from zebrafish demonstrated that it is inactive with perfectly complementary targets but a mismatch in the seed region stimulates target RNA cleavage (44). It can be proposed that mismatches in the seed region may allosterically change the active site conformation or target positioning in the case of zAgo2 or LrAgo. In contrast, target DNA cleavage by CbAgo and LrAgo is strongly inhibited in the presence of mismatches in the 3’-supplementary guide region. Therefore, in the case of these pAgos propagation of the guide-target duplex, rather than initial interactions in the seed region, may control the fidelity of target recognition.

Both CbAgo and LrAgo can relax supercoiled plasmid DNA in a guide-independent manner, likely by accommodating both DNA strands within the catalytic cleft, thus demonstrating the so-called ‘chopping’ activity previously described for TtAgo and MjAgo (11,27). Plasmid relaxation is followed by slow processing of plasmid DNA, resulting in its linearization and further degradation. The chopping activity may complicate the use of pAgos as specific genome editing tools, as recently discussed for TtAgo and NgAgo (30). Indeed, LrAgo can process plasmid DNA with similar efficiency independently of guide binding. In contrast, the chopping activity of CbAgo is markedly suppressed when it is preloaded with guide molecules. Furthermore, CbAgo can precisely cut dsDNA at one or more sites when programmed with corresponding guides. This opens the way for development of novel pAgo-based tools for DNA manipulations *in vitro* and *in vivo*.

Previous attempts to use thermophilic pAgos, such as PfAgo or TtAgo, as programmable nucleases were limited by their low activity at ambient temperature, which required heating the samples with concomitant DNA denaturation (13,28). We showed that CbAgo can target specific DNA sites at much lower temperatures, with low efficiency at 37°C and with high efficiency at 55°C - the temperature compatible with many *in vitro* applications. One obvious application is the use CbAgo in recombinant DNA technology, by analogy with restriction endonucleases but with potential ability to target any site of interest. In contrast to restriction endonucleases and Cas9, guide-directed CbAgo cuts only one DNA strand in the dsDNA duplex. Therefore, two strands of DNA can be targeted by CbAgo loaded with two guides independently, so ‘sticky’ ends of any desired configuration can be produced. In contrast to Cas nucleases, CbAgo do not require the presence of any specific motifs (such as PAM, protospacer adjacent motif) in the guide or target DNAs which may enable DNA targeting with a single-nucleotide resolution. Furthermore, short DNA oligonucleotides utilized by CbAgo as guide molecules are much easier to synthesize compared to longer RNA guides required for Cas nucleases.

Important problems that need to be solved to allow the use of pAgos in genetic technologies such as genome editing include finding of conditions that would allow specific guide loading and efficient targeting of dsDNA *in vivo* in eukaryotic cells. In contrast to Cascade complexes and Cas nucleases of the CRISPR systems, pAgos do not unwind dsDNA so they might interact with auxiliary cellular factors that could perform DNA melting or recruit pAgos to ssDNA regions. Indeed, DNA cleavage by TtAgo was recently shown to be stimulated by SSB or UvrD helicase (45). Furthermore, negative DNA supercoiling might facilitate the formation of ternary pAgo complexes at locally unwound plasmid or genomic strands (8,13). Analysis of *in vivo* DNA processing by mesophilic pAgos and its dependence on various cellular activities will be an important goal of future studies.

## ACKNOWLEDGMENTS

We thank D. Esyunina and M. Petrova for experimental support and helpful discussions, A. Oguienko for assistance with figure preparation.

## FUNDING

This work was supported by the Grant of the Ministry of Higher Education and Science of Russian Federation 14.W03.31.0007. The authors declare no competing interests.

## Supplementary Information

**Figure S1.**
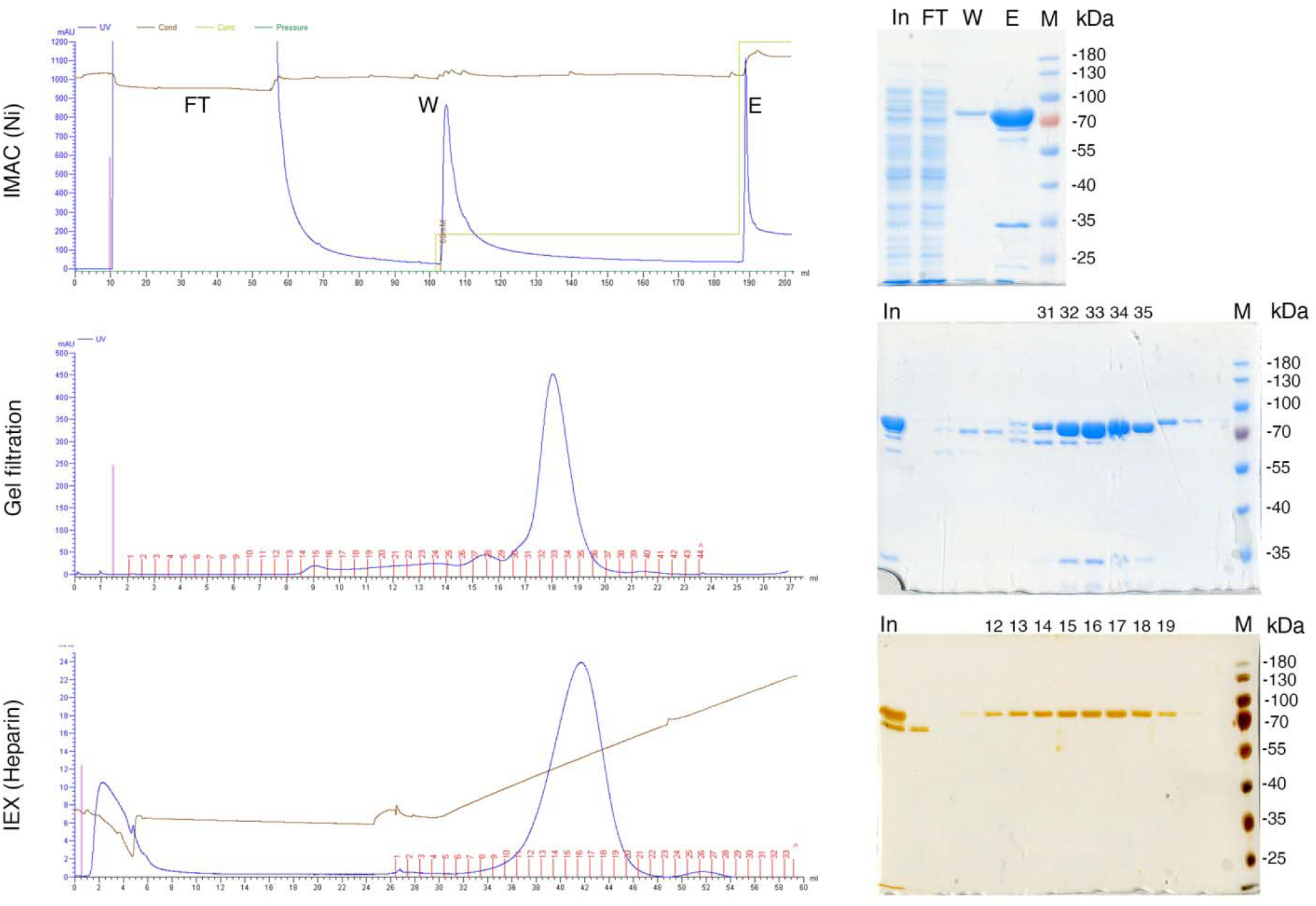
Three-step purification of LrAgo. (*Upper panel*) Ni-NTA-column chromatogram (left) and a representative SDS-PAGE gel of indicated fractions (In, input, FT, flowthrough; W, 60 mM imidazole wash; E, 270 mM imidazole elution; M, marker). (*Middle panel*) Gel filtration chromatogram (left) and a representative gel of indicated fractions (In, input; 31-35, chromatography fractions containing LrAgo). (*Lower panel*) Chromatography on heparin column (left) and a representative gel of indicated fractions (silver staining).

**Figure S2.**
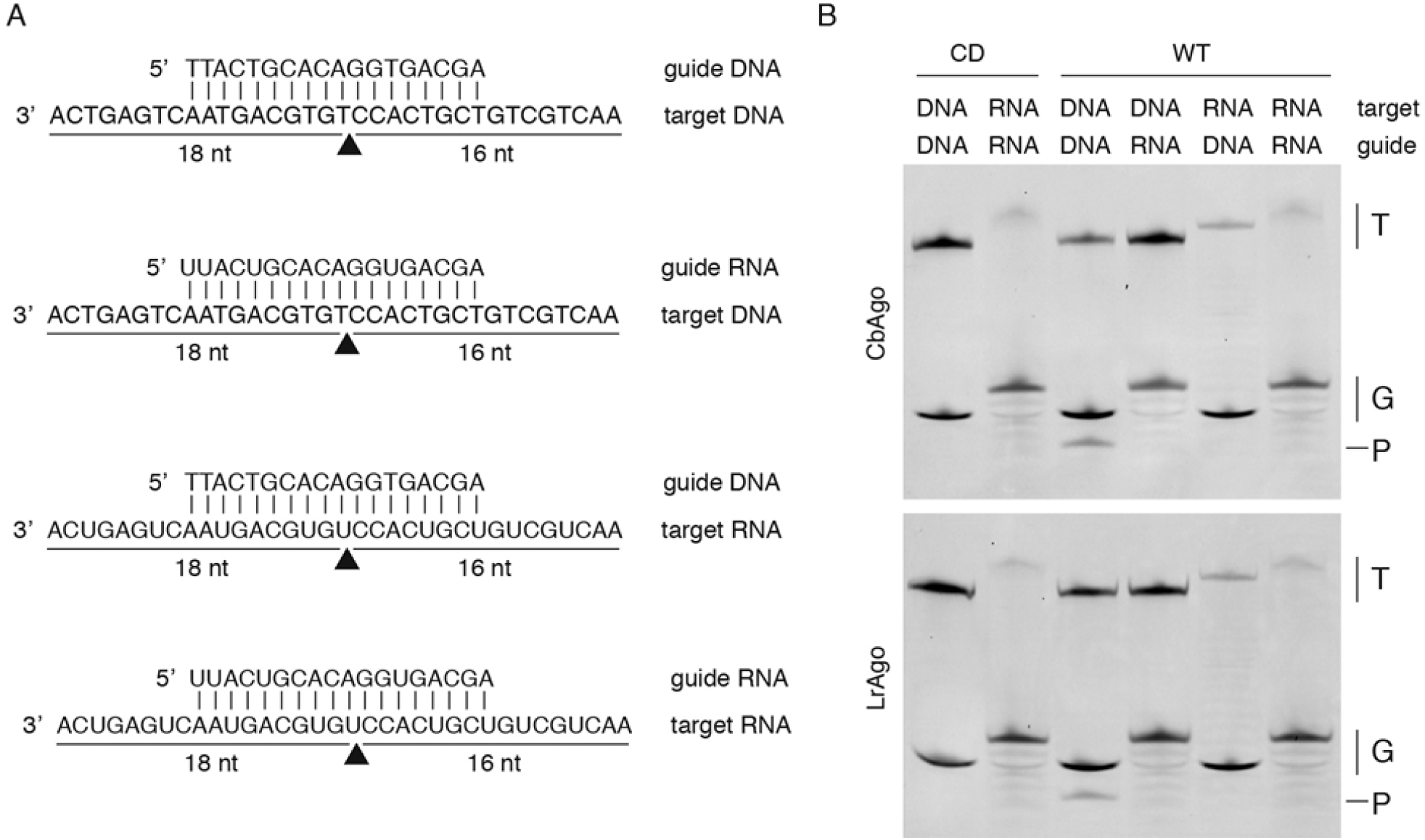
(A) DNA and RNA oligonucleotides used for *in vitro* analysis of the guide/target specificity of pAgos. Black triangle indicates the predicted cleavage site; the lengths of the cleavage products are indicated. (B) The cleavage assay with DNA or RNA guide/target oligonucleotides. Both CbAgo and LrAgo can cleave ssDNA but not ssRNA when loaded with complementary DNA guide. Note that the size of one of the cleavage products coincides with the size of the guide DNA. RNA guides are not able to direct efficient slicing of either complementary RNA or DNA targets. T, target; P, product; G, guide; WT, wild-type pAgo; CD, catalytically dead pAgo.

**Figure S3.**
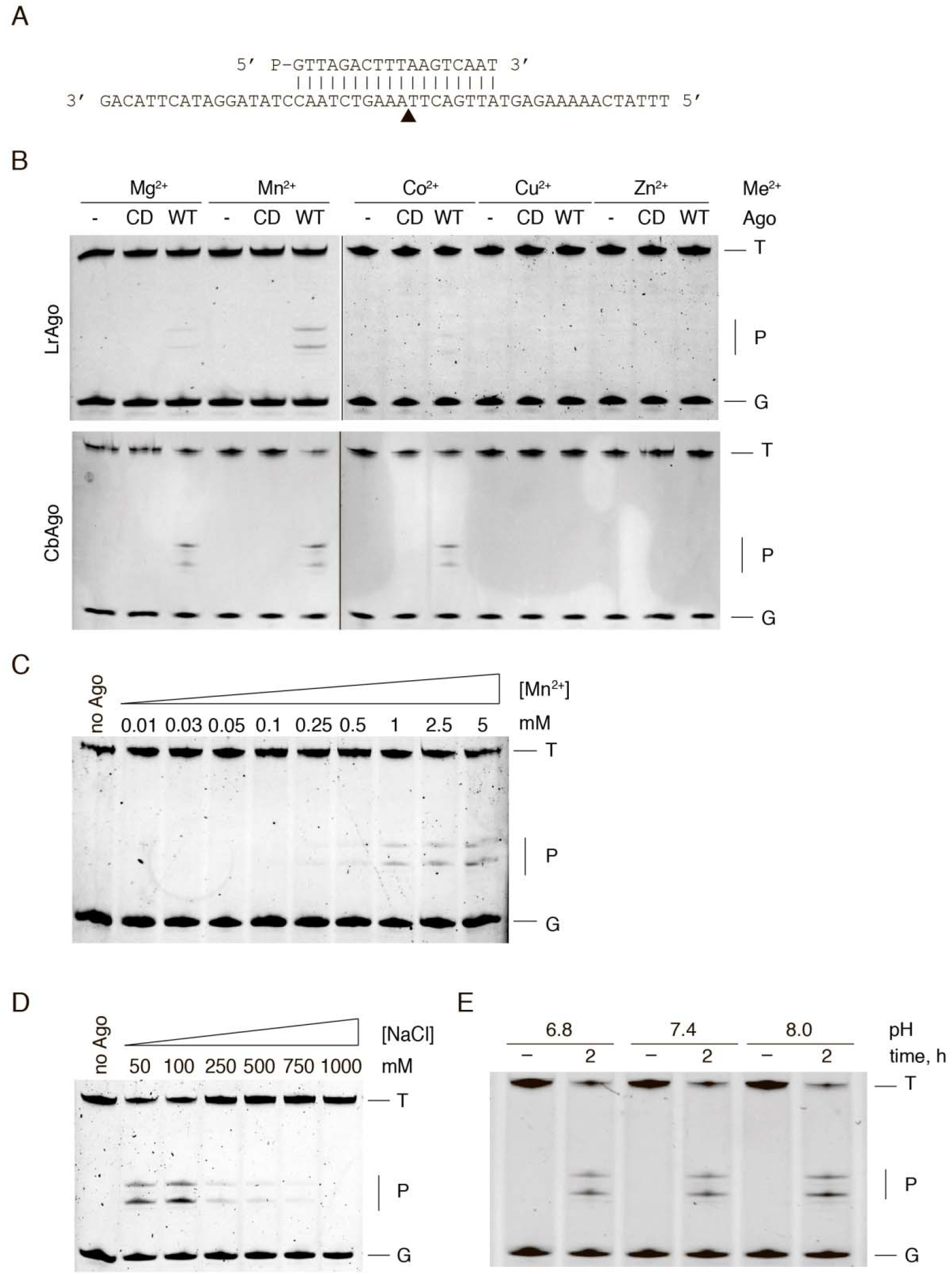
Biochemical properties of LrAgo and CbAgo. (A) Guide and target DNA oligonucleotides used for *in vitro* assays. Black triangle indicates the predicted cleavage site. Effects of (B) different cations, (C) Mn^2+^ concentration, (D) NaCl concentration, and (E) pH on pAgo activity. All reactions were carried out for 2 hours at 37 °C, at the 5:5:1 pAgo:guide:target molar ratio. Positions of the guide (G), target (T) and cleavage products (P) are indicated.

**Figure S4.**
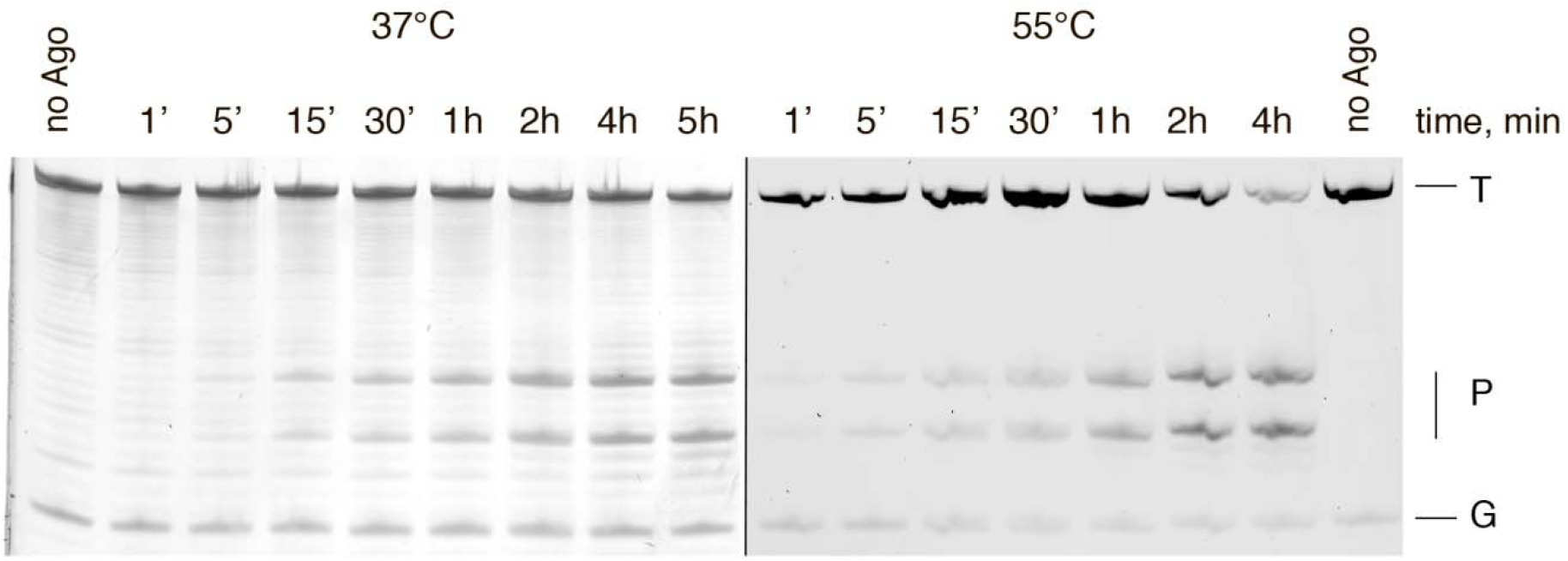
Kinetics of target cleavage by CbAgo at 37 and 55 °C. CbAgo was mixed with target DNA after preliminary loading with complementary guide at the 5:1:5 pAgo:guide:target molar ratio. Under these conditions, the DNA target was in a 5-fold molar excess over the binary pAgo-guide complex. The samples were incubated at either 37 or 55 °C for indicated time intervals; the reaction products were resolved by 19% urea-PAGE and stained with SYBR Gold. At 37°C (left panel), the cleavage does not go to completion after reaching a certain level, suggesting that the reaction has a limited turnover possibly due to slow dissociation of the product complex. At 55°C (right panel), the efficiency of cleavage is increased, multi-turnover catalysis under these conditions. T, target DNA; G, guide DNA; P, DNA products.

**Figure S5.**
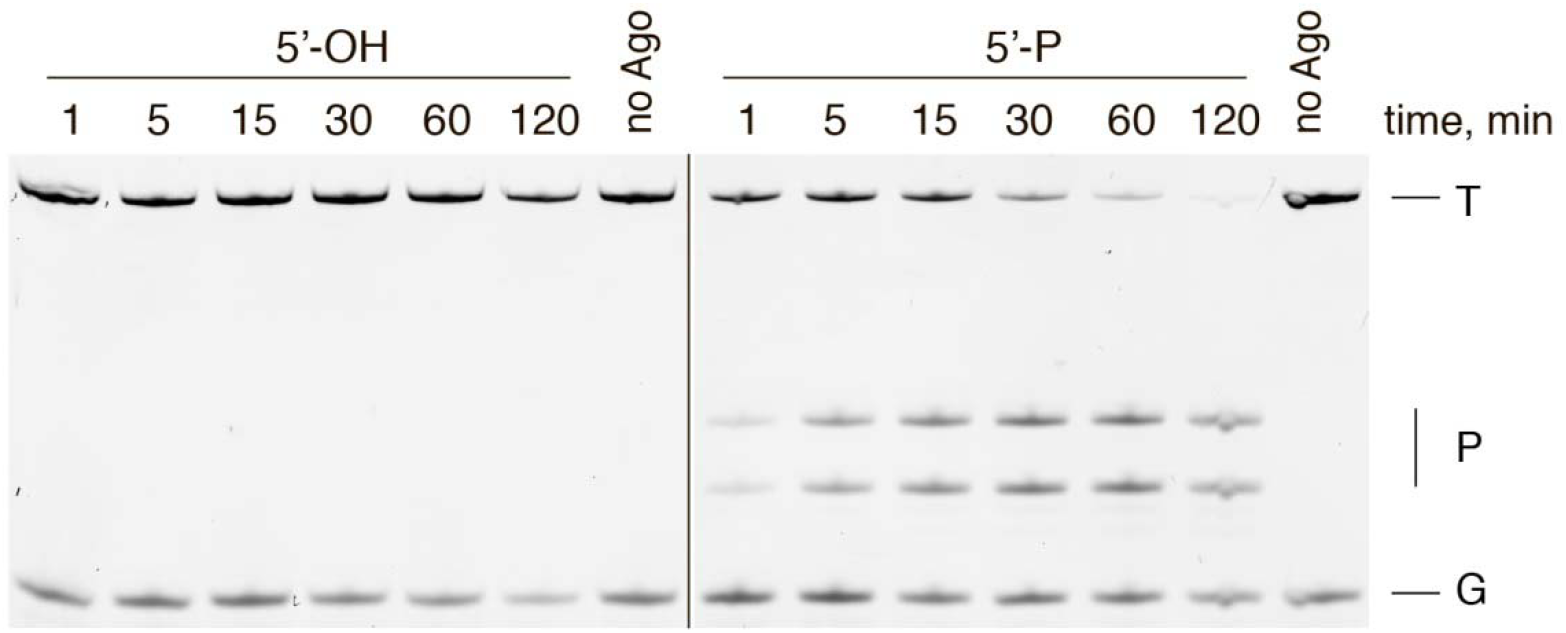
Analysis of ssDNA cleavage by CbAgo loaded with complementary 5’-OH and 5’-P guides at 55°C. The reactions were performed at the 5:2:1 pAgo:guide:target molar ratio for indicated time periods. The absence of activity in reactions with 5’-OH guides suggests the inability of CbAgo to utilize nonphosphorylated DNA guides at 55 °C.

**Figure S6.**
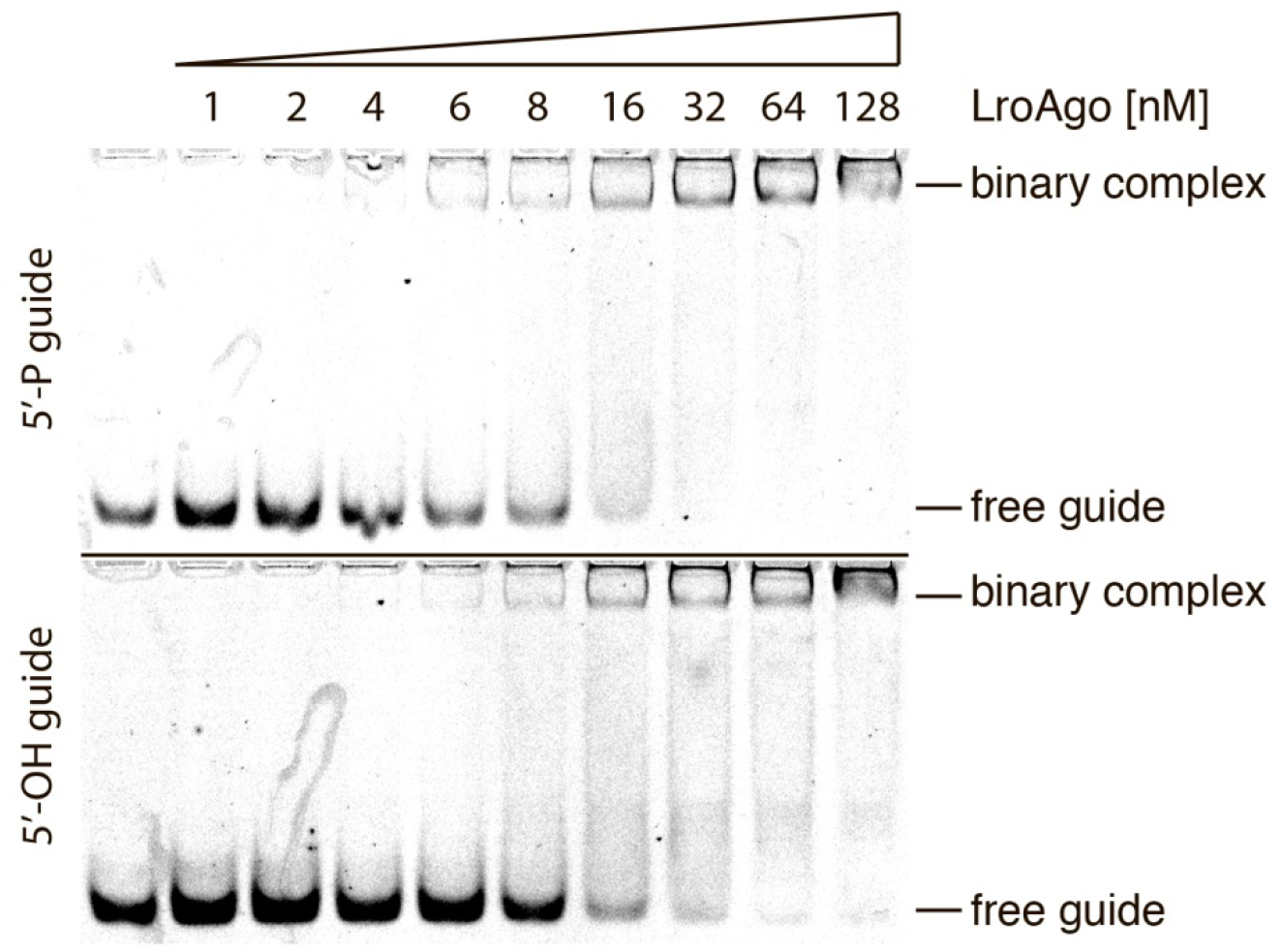
LrAgo forms binary complexes with 5’-P and 5’-OH guides with the same efficiency. Guide DNA (5 nM) was incubated with increasing amounts of LrAgo at 37°C for 10 minutes and resolved by native 10% PAGE. Positions of the free guide as well as the binary complex are indicated. EMSA was conducted in quadruplicate and a representative gel is shown.

**Figure S7.**
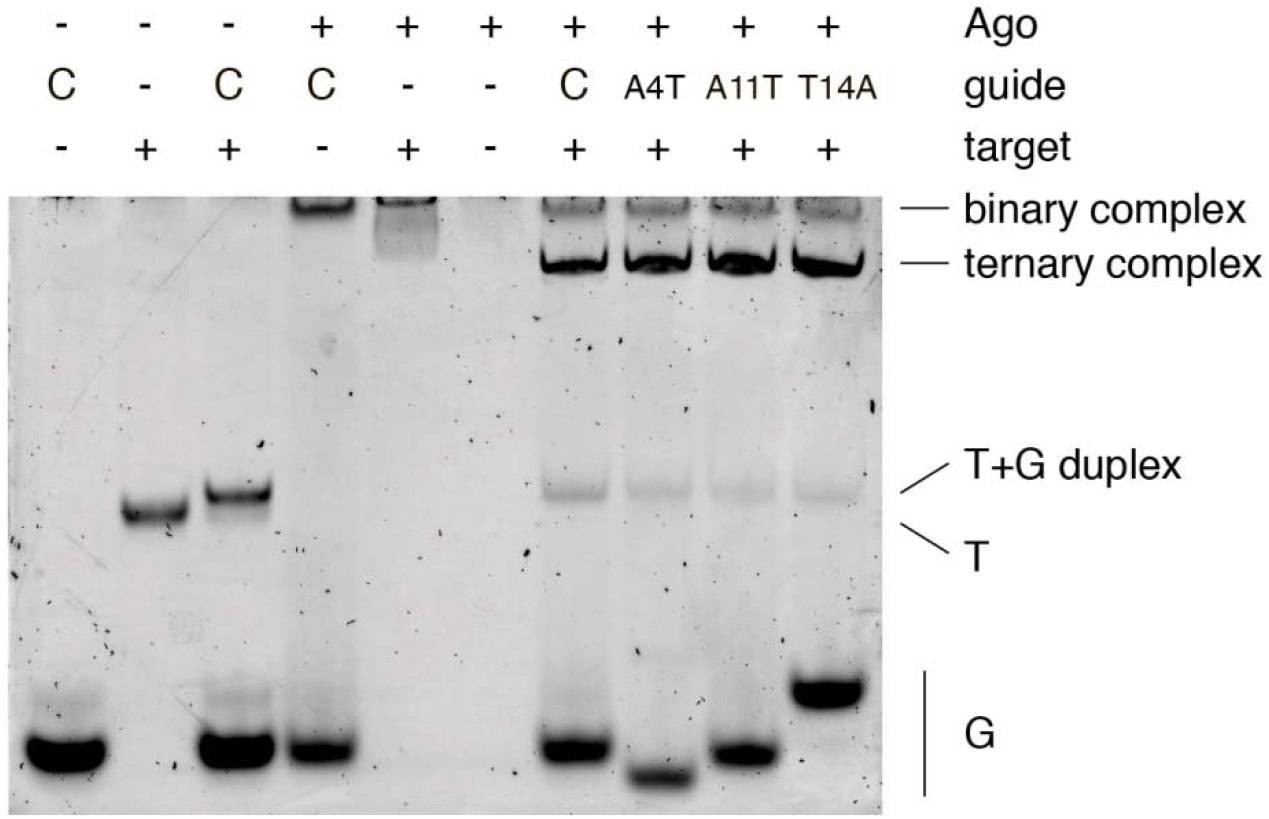
Formation of binary and ternary complexes with guide and target DNA by LrAgo. The complexes were formed for 10 minutes at 37 °C and resolved by 10% native PAGE. Mismatches in guide DNA that significantly alter the efficiency of LrAgo-mediated cleavage (Fig. 4) do not affect target binding at the given time-scale. C, fully complementary guide DNA; T, target DNA; G, guide DNA; nucleotide substitutions in the mismatched guides are designated above the gel.

**Table S1.**
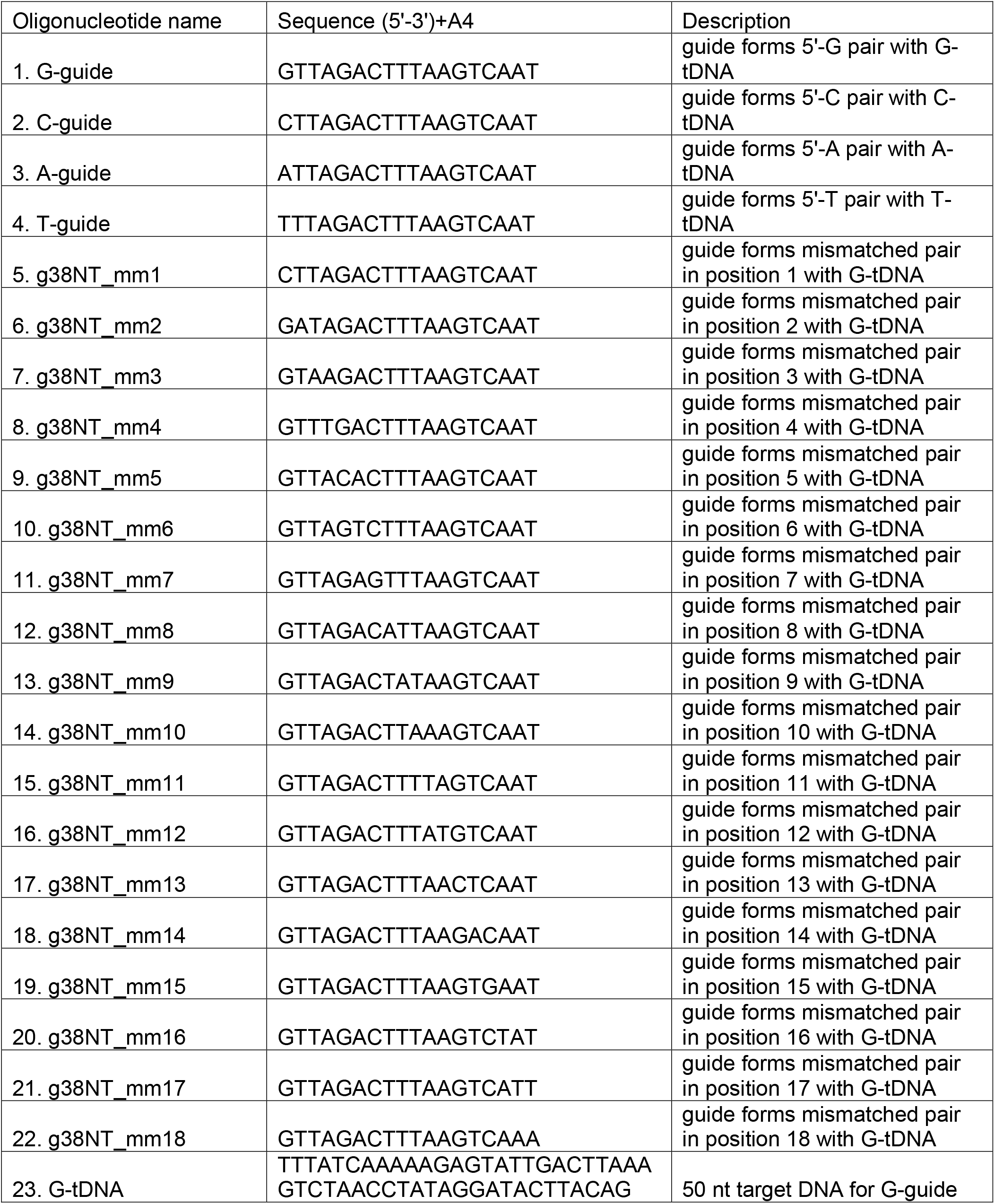
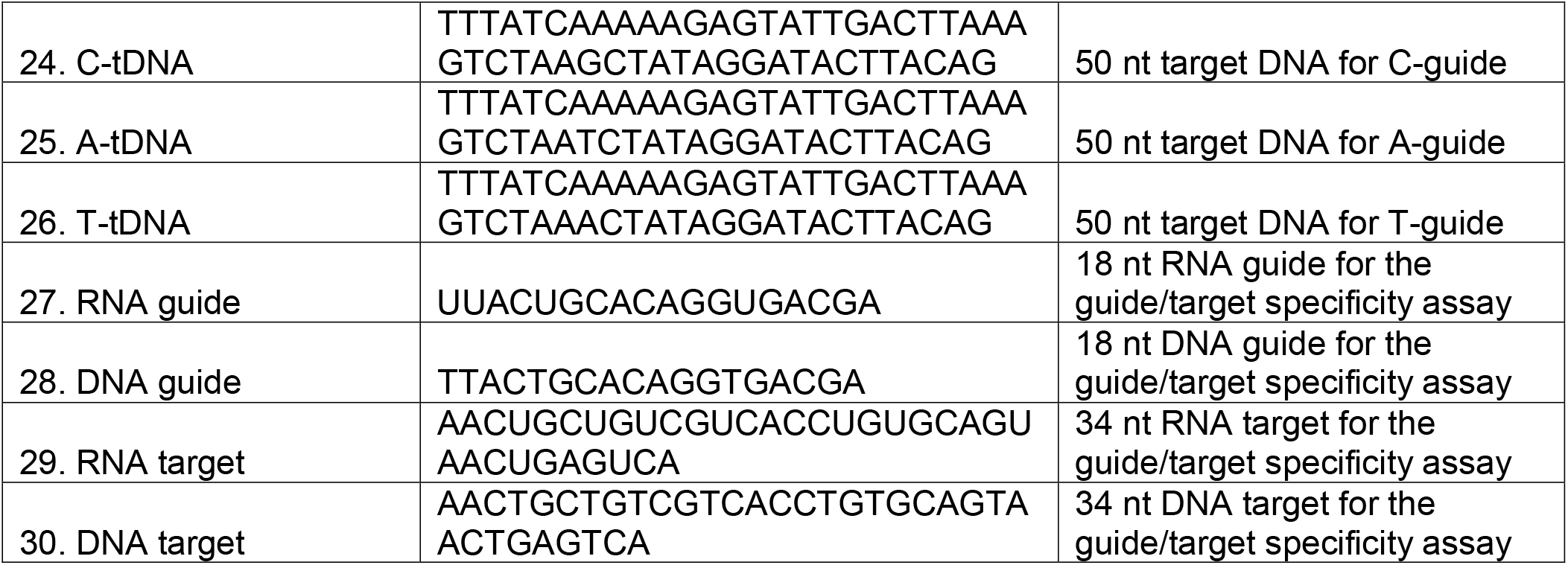
Sequences of oligonucleotides used in the cleavage assays.

## REFERENCES

1. Joshua-Tor, L. (2006) The Argonautes. Cold Spring Harbor symposia on quantitative biology, 71, 67–72.

2. Peters, L. and Meister, G. (2007) Argonaute proteins: mediators of RNA silencing. Molecular cell, 26, 611–623.

3. Pratt, A.J. and MacRae, I.J. (2009) The RNA-induced silencing complex: a versatile gene-silencing machine. The Journal of biological chemistry, 284, 17897–17901.

4. Ryazansky, S., Kulbachinskiy, A. and Aravin, A.A. (2018) The Expanded Universe of Prokaryotic Argonaute Proteins. mBio, 9.

5. Makarova, K.S., Wolf, Y.I., van der Oost, J. and Koonin, E.V. (2009) Prokaryotic homologs of Argonaute proteins are predicted to function as key components of a novel system of defense against mobile genetic elements. Biology direct, 4, 29.

6. Swarts, D.C., Makarova, K., Wang, Y., Nakanishi, K., Ketting, R.F., Koonin, E.V., Patel, D.J. and van der Oost, J. (2014) The evolutionary journey of Argonaute proteins. Nature structural & molecular biology, 21, 743–753.

7. Kaya, E., Doxzen, K.W., Knoll, K.R., Wilson, R.C., Strutt, S.C., Kranzusch, P.J. and Doudna, J.A. (2016) A bacterial Argonaute with noncanonical guide RNA specificity. Proceedings of the National Academy of Sciences of the United States of America, 113, 4057–4062.

8. Swarts, D.C., Hegge, J.W., Hinojo, I., Shiimori, M., Ellis, M.A., Dumrongkulraksa, J., Terns, R.M., Terns, M.P. and van der Oost, J. (2015) Argonaute of the archaeon Pyrococcus furiosus is a DNA-guided nuclease that targets cognate DNA. Nucleic acids research, 43, 5120–5129.

9. Wang, Y., Juranek, S., Li, H., Sheng, G., Tuschl, T. and Patel, D.J. (2008) Structure of an argonaute silencing complex with a seed-containing guide DNA and target RNA duplex. Nature, 456, 921–926.

10. Yuan, Y.R., Pei, Y., Ma, J.B., Kuryavyi, V., Zhadina, M., Meister, G., Chen, H.Y., Dauter, Z., Tuschl, T. and Patel, D.J. (2005) Crystal structure of A. aeolicus argonaute, a site-specific DNA-guided endoribonuclease, provides insights into RISC-mediated mRNA cleavage. Molecular cell, 19, 405–419.

11. Zander, A., Willkomm, S., Ofer, S., van Wolferen, M., Egert, L., Buchmeier, S., Stockl, S., Tinnefeld, P., Schneider, S., Klingl, A. et al. (2017) Guide-independent DNA cleavage by archaeal Argonaute from Methanocaldococcus jannaschii. Nature microbiology, 2, 17034.

12. Olovnikov, I., Chan, K., Sachidanandam, R., Newman, D.K. and Aravin, A.A. (2013) Bacterial argonaute samples the transcriptome to identify foreign DNA. Molecular cell, 51, 594–605.

13. Swarts, D.C., Jore, M.M., Westra, E.R., Zhu, Y., Janssen, J.H., Snijders, A.P., Wang, Y., Patel, D.J., Berenguer, J., Brouns, S.J.J. et al. (2014) DNA-guided DNA interference by a prokaryotic Argonaute. Nature, 507, 258–261.

14. Lisitskaya, L., Aravin, A.A. and Kulbachinskiy, A. (2018) RNA interference and beyond: structure and functions of prokaryotic Argonaute proteins. Nature communications, 9, 5165.

15. Koonin, E.V. (2017) Evolution of RNA- and DNA-guided antivirus defense systems in prokaryotes and eukaryotes: common ancestry vs convergence. Biology direct, 12, 5.

16. Hegge, J.W., Swarts, D.C. and van der Oost, J. (2017) Prokaryotic Argonaute proteins: novel genome-editing tools? Nature reviews. Microbiology, 16, 5–11.

17. Liu, Y., Esyunina, D., Olovnikov, I., Teplova, M., Kulbachinskiy, A., Aravin, A.A. and Patel, D.J. (2018) Accommodation of helical imperfections in *Rhodobacter sphaeroides* Argonaute ternary complexes with guide RNA and target DNA. Cell reports, 24, 453–462.

18. Miyoshi, T., Ito, K., Murakami, R. and Uchiumi, T. (2016) Structural basis for the recognition of guide RNA and target DNA heteroduplex by Argonaute. Nature communications, 7, 11846.

19. Sheng, G., Zhao, H., Wang, J., Rao, Y., Tian, W., Swarts, D.C., van der Oost, J., Patel, D.J. and Wang, Y. (2014) Structure-based cleavage mechanism of Thermus thermophilus Argonaute DNA guide strand-mediated DNA target cleavage. Proceedings of the National Academy of Sciences of the United States of America, 111, 652–657.

20. Wang, Y., Sheng, G., Juranek, S., Tuschl, T. and Patel, D.J. (2008) Structure of the guide-strand-containing argonaute silencing complex. Nature, 456, 209–213.

21. Willkomm, S., Oellig, C.A., Zander, A., Restle, T., Keegan, R., Grohmann, D. and Schneider, S. (2017) Structural and mechanistic insights into an archaeal DNA-guided Argonaute protein. Nature microbiology, 2, 17035.

22. Parker, J.S., Parizotto, E.A., Wang, M., Roe, S.M. and Barford, D. (2009) Enhancement of the seed-target recognition step in RNA silencing by a PIWI/MID domain protein. Molecular cell, 33, 204–214.

23. Zander, A., Holzmeister, P., Klose, D., Tinnefeld, P. and Grohmann, D. (2014) Single-molecule FRET supports the two-state model of Argonaute action. RNA biology, 11, 45–56.

24. Salomon, W.E., Jolly, S.M., Moore, M.J., Zamore, P.D. and Serebrov, V. (2015) Single-Molecule Imaging Reveals that Argonaute Reshapes the Binding Properties of Its Nucleic Acid Guides. Cell, 162, 84–95.

25. Wee, L.M., Flores-Jasso, C.F., Salomon, W.E. and Zamore, P.D. (2012) Argonaute divides its RNA guide into domains with distinct functions and RNA-binding properties. Cell, 151, 1055–1067.

26. Doxzen, K.W. and Doudna, J.A. (2017) DNA recognition by an RNA-guided bacterial Argonaute. PloS one, 12, e0177097.

27. Swarts, D.C., Szczepaniak, M., Sheng, G., Chandradoss, S.D., Zhu, Y., Timmers, E.M., Zhang, Y., Zhao, H., Lou, J., Wang, Y. et al. (2017) Autonomous Generation and Loading of DNA Guides by Bacterial Argonaute. Molecular cell, 65, 985–998.

28. Enghiad, B. and Zhao, H. (2017) Programmable DNA-Guided Artificial Restriction Enzymes. ACS synthetic biology, 6, 752–757.

29. Lee SH T.G., Ata H, Nowsheen S, Romito M, Lou Z, Ryu SM, Ekker SC, Cathomen T, Kim JS. (2017) Failure to detect DNA-guided genome editing using *Natronobacterium gregoryi* Argonaute. Nature biotechnology, 35, 17–18.

30. O’Geen, H., Ren, C., Coggins, N.B., Bates, S.L. and Segal, D.J. (2018) Unexpected binding behaviors of bacterial Argonautes in human cells cast doubts on their use as targetable gene regulators. PloS one, 13, e0193818.

31. Oganesyan, N., Ankoudinova, I., Kim, S.H. and Kim, R. (2007) Effect of osmotic stress and heat shock in recombinant protein overexpression and crystallization. Protein expression and purification, 52, 280–285.

32. Wong, I. and Lohman, T.M. (1993) A double-filter method for nitrocellulose-filter binding: application to protein-nucleic acid interactions. Proceedings of the National Academy of Sciences of the United States of America, 90, 5428–5432.

33. Keren, N., Kidd, M.J., Penner-Hahn, J.E. and Pakrasi, H.B. (2002) A light-dependent mechanism for massive accumulation of manganese in the photosynthetic bacterium Synechocystis sp. PCC 6803. Biochemistry, 41, 15085–15092.

34. Ghildiyal, M. and Zamore, P.D. (2009) Small silencing RNAs: an expanding universe. Nat Rev Genet, 10, 94–108.

35. Elkayam, E., Kuhn, C.D., Tocilj, A., Haase, A.D., Greene, E.M., Hannon, G.J. and Joshua-Tor, L. (2012) The structure of human argonaute-2 in complex with miR-20a. Cell, 150, 100–110.

36. Matsumoto, N., Nishimasu, H., Sakakibara, K., Nishida, K.M., Hirano, T., Ishitani, R., Siomi, H., Siomi, M.C. and Nureki, O. (2016) Crystal Structure of Silkworm PIWI-Clade Argonaute Siwi Bound to piRNA. Cell, 167, 484–497.

37. Schirle, N.T. and MacRae, I.J. (2012) The crystal structure of human Argonaute2. Science, 336, 1037–1040.

38. Dahlgren, C., Zhang, H.Y., Du, Q., Grahn, M., Norstedt, G., Wahlestedt, C. and Liang, Z. (2008) Analysis of siRNA specificity on targets with double-nucleotide mismatches. Nucleic acids research, 36, e53.

39. Doench, J.G. and Sharp, P.A. (2004) Specificity of microRNA target selection in translational repression. Genes & development, 18, 504–511.

40. Sun, G., Wang, J., Huang, Y., Yuan, C.W., Zhang, K., Hu, S., Chen, L., Lin, R.J., Yen, Y. and Riggs, A.D. (2018) Differences in silencing of mismatched targets by sliced versus diced siRNAs. Nucleic acids research, 46, 6806–6822.

41. Sheng, G., Gogakos, T., Wang, J., Zhao, H., Serganov, A., Juranek, S., Tuschl, T., Patel, D.J. and Wang, Y. (2017) Structure/cleavage-based insights into helical perturbations at bulge sites within T. thermophilus Argonaute silencing complexes. Nucleic acids research, 45, 9149–9163.

42. Willkomm, S., Zander, A., Grohmann, D. and Restle, T. (2016) Mechanistic Insights into Archaeal and Human Argonaute Substrate Binding and Cleavage Properties. PloS one, 11, e0164695.

43. Rivas, F.V., Tolia, N.H., Song, J.J., Aragon, J.P., Liu, J., Hannon, G.J. and Joshua-Tor, L. (2005) Purified Argonaute2 and an siRNA form recombinant human RISC. Nature structural & molecular biology, 12, 340–349.

44. Chen, G.R., Sive, H. and Bartel, D.P. (2017) A Seed Mismatch Enhances Argonaute2-Catalyzed Cleavage and Partially Rescues Severely Impaired Cleavage Found in Fish. Molecular cell, 68, 1095–1107.

45. Hunt, E.A., Evans, T.C., Jr. and Tanner, N.A. (2018) Single-stranded binding proteins and helicase enhance the activity of prokaryotic argonautes in vitro. PloS one, 13, e0203073.

